# Piezo1–Yap1 signalling translates tissue mechanics into regenerative cell states

**DOI:** 10.1101/2025.09.20.677431

**Authors:** Fernando Ferreira, Jaime A. Espina, Artemis G. Korovesi, Inês A. Ferreira, Quentin Tirel, Elias H. Barriga

## Abstract

Epimorphic regeneration encompasses several stages, such as wound healing, formation of a regenerative bud and regenerative outgrowth. The signatures that define each stage have been well studied, yet little is known about the mechanisms that coordinate cell state transitions across regenerative stages. Here, we show that stiffening of wounded epithelial tissues is essential for the formation of a prospective regenerative bud and, in turn, for the transition from wound healing to bud formation. Mechanistically, to detect prospective bud stiffening, cells use a mechanosensitive cascade involving Piezo1-dependent Yap1 activation. We also determined that this cascade is required for the acquisition of a transcriptional profile that endows bud cells with regenerative competence. Notably, the activation of this Piezo1-Yap1 mechanosensitive cascade in nonregenerative contexts is sufficient to induce the formation of competent bud-like structures, which are sufficient to drive *de novo* regeneration in these otherwise incompetent tissues. Taken together, our results reveal a mechanical state at the boundary between wound healing and regenerative bud formation, which is essential for synchronizing the transition between these pivotal stages of regeneration. More broadly, these findings support the general idea that the physical properties of tissues integrate with molecular frameworks to actuate cell state transitions during morphogenesis.

## Main

Some vertebrates, particularly amphibians and lizards, can regenerate lost appendages via a process known as epimorphic regeneration or epimorphosis^1^. During this process, cells within regenerating tissues synchronize their activities to transit across multiple regenerative stages^2^, such as wound healing, blastema or regenerative bud formation, and regenerative outgrowth^3,4^. The cellular and biochemical signatures that define these phases have been extensively studied^5–7^, and more recently, these approaches have been complemented by biophysical profiling^8–13^. However, in addition to their wide relevance for our understanding of regenerative processes, the mechanisms that coordinate biochemical signalling with cell behaviours to synchronize cell state transitions across regenerative stages remain less understood. Mechanical cues contribute to synchronizing cell state transitions during embryonic development^14–17^, a context that involves similar events of tissue remodelling and signalling cascades to those operating during regeneration^18,19^. Moreover, the regeneration of an appendage is inherently influenced by mechanical forces, and mechanosensitive molecules are involved in this process^20^. Thus, we hypothesized that mechanical signals arising from the regenerative microenvironment are part of the mechanism that instructs cells within wounded tissues to transit across regenerative states.

To study the contribution of tissue mechanics to cell state transitions during regeneration, we used *Xenopus laevis* larval tails, a well-established model of vertebrate epimorphic regeneration (**Fig. 1a**)^21–24^. During an amputation assay (**Suppl Fig. 1a–c**; details in **Methods**), the regeneration of *Xenopus* larval tails involves 3 major phases: wound healing, in which a specialized wound epithelia closes the wound; regenerative bud formation, a hallmark of regeneration as the bud is a blastema-like mass of undifferentiated cells from where tissues regenerate; and regenerative outgrowth, in which tissues grow from the bud until a full tail is regenerated (**Fig. 1a**)^3,21^. While the cellular and biochemical events that define the stages of *Xenopus* tail regeneration have been well studied^3,21^, relatively little is known about their mechanical signatures^25^. Thus, to explore the relevance of tissue mechanics for cell state transitions during regeneration, we temporally mapped the apparent elastic moduli (hereafter referred to as stiffness) of regenerating tissues via *in vivo* atomic force microscopy (*i*AFM) (**Fig. 1b**; **Suppl Fig. 2a, b;** details in **Methods**). Owing to the relevance of the regenerative bud for the initiation and patterning of tail regeneration^23,24^, we directly measured prospective bud tissues (**Fig. 1c**). *i*AFM analyses revealed a steep increase in tissue stiffness at early stages of wound healing (from 1 to 6 hours post amputation, hpa); this stiffening plateaued once a prospective regenerative bud (6 to 24 hpa), and the stiffness values remained stable at later time points, which are relevant for subsequent transitions (24 and 48 hpa) (**Fig. 1d**). The observed increase in tissue stiffening temporally correlates with the establishment of prospective bud tissues (**Fig. 1e**), suggesting that tissue stiffening may instruct cells within wounded epithelia to transit into bud-like regenerative states.

**Fig. 1.**
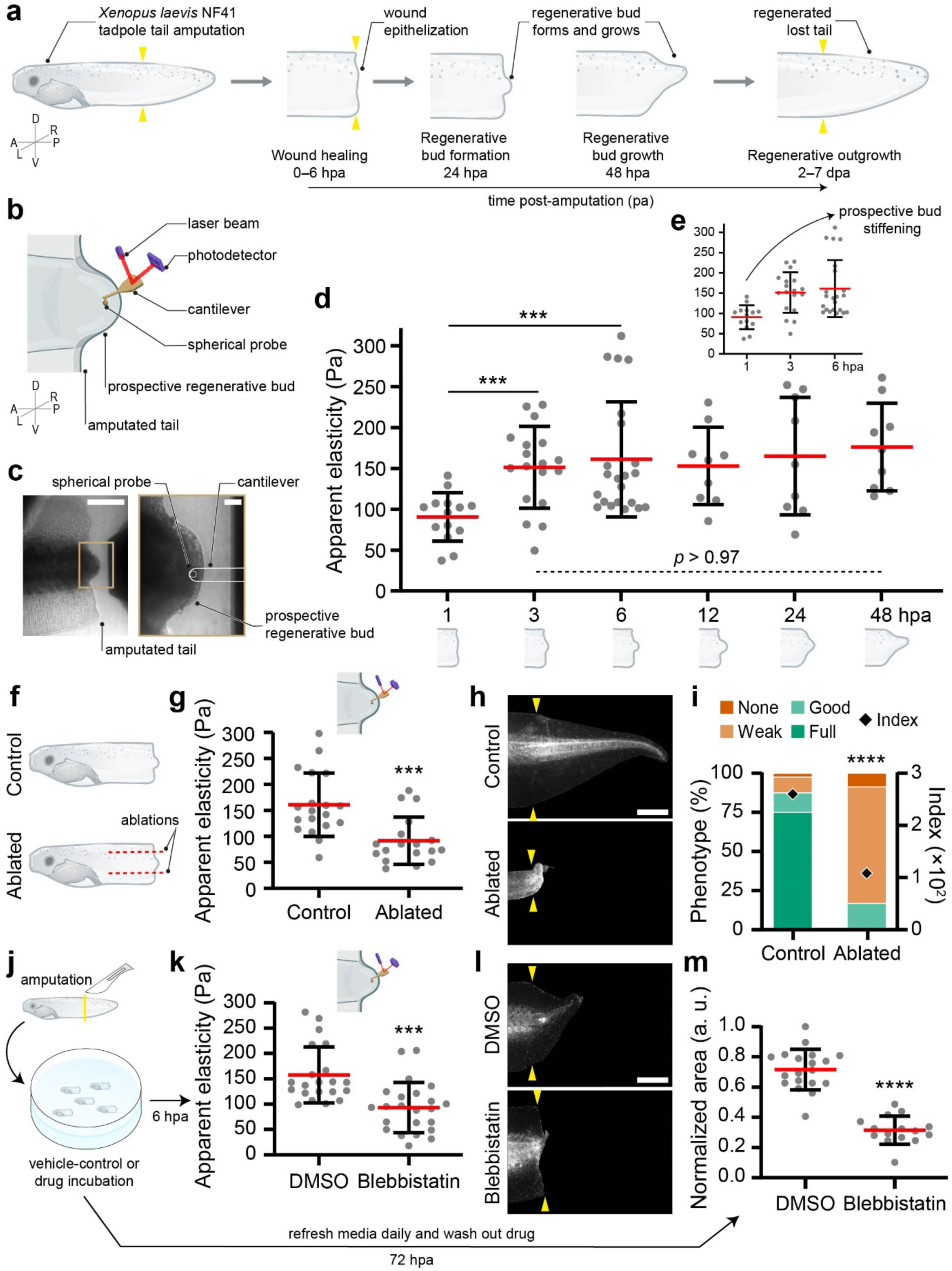
Tissue stiffening at the transition from wound healing to regenerative bud formation. **a**, Scheme of the phases and transitions of regeneration. A, anterior, P, posterior; D, dorsal; V, ventral; L, left; R, right. Yellow arrowheads, amputation plane. **b**, Scheme of *i*AFM measurements. **c**, Cantilever positioning relative to prospective regenerative bud at 6 hpa (left panel). Zoom-in with cantilever and probe outlined in white line (right panel). Scale bars, 500 µm (left panel) and 50 µm (right panel). **d**, Tissue stiffening measurements during 48 hpa. **e**, Detail of the time-points depicting the prospective bud stiffening at the transition. Red lines represent mean and error bars standard deviation. Two-tailed Student’s *t*-test, \*\*\**p*_1 vs. 3 hpa_ = 0.0004, \*\*\**p*_1 vs. 6 hpa_ = 0.0002 (with Welch’s correction), Brown-Forsythe and Welch ANOVA with Dunnett’s multiple comparisons test, *p* > 0.97 for all comparisons between 3 and 48 hpa, *n*_1 hpa_ = 14, *n*_3 hpa_ = 18, *n*_6 hpa_ = 22, *n*_12 hpa_ = 9, *n*_24 hpa_ = 9, *n*_48 hpa_ = 9 tails. **f–i**, Tissue ablation decreases tissue stiffening that is required for regeneration. **f**, Scheme of tissue ablation experiment. **g**, Quantification of tissue stiffening at 6 hpa. Red lines represent mean and error bars standard deviation. Two-tailed Student’s *t*-test, \*\*\**p* = 0.0004, *n*_Control_ = 19, *n*_Ablated_ = 18 tails. **h**, Tails at 7 dpa. Scale bar, 500 µm. **i**, Quantification of regeneration efficiency. Two-tailed Fisher’s exact test, \*\*\*\**p* < 0.0001, *n*_Control_ = 33, *n*_Ablated_ = 26 tails. **j–m**, Blebbistatin decreases tissue stiffening that is required for regeneration. **j**, Scheme of regeneration assay. **k**, Quantification of tissue stiffening. Incubation with 0.1% DMSO or 50 µM blebbistatin from 1 to 6 hpa. Red lines represent mean and error bars standard deviation. Two-tailed Student’s *t-*test, \*\*\**p* = 0.0002, *n*_DMSO_ = 21, *n*_Blebbistatin_ = 23 tails. **l**, Tails at 72 hpa. Incubation with 0.1% DMSO or 50 µM blebbistatin from 1 to 24 hpa. Scale bar, 500 µm. **m**, Quantification of regeneration efficiency. Red lines represent mean and error bars standard deviation. Two-tailed Student’s *t-*test, \*\*\*\**p* < 0.0001, *n*_DMSO_ = 19, *n*_Blebbistatin_ = 15 tails. **c,h,l**, Representative examples from at least three independent experiments; CI = 95%.

To test whether early stiffening of prospective bud tissues is relevant for regeneration, we prevented this mechanical build-up by using mechanical and molecular perturbations. Most living systems are under tension, which often leads to strain stiffening of tissues, increasing their apparent elastic moduli^14,26–31^. Thus, to prevent prospective bud stiffening, we adapted a previously used method that relies on oriented mechanical ablations that release tension and, in turn, prevent tissue stiffening^14,23^. Strain of developing *Xenopus laevis* posterior epithelial cells has been reported^32^, and our morphological analyses revealed a dorsoventral distribution of membrane stress in the tissues surrounding the prospective bud (**Suppl Fig. 2c**). Therefore, ablations were performed from anterior to posterior, perpendicularly to the dorsoventral axis (**Fig. 1f**; details in **Methods**). *i*AFM measurements confirmed that oriented ablation was sufficient to prevent stiffening of the prospective bud (**Fig. 1f, g**). Notably, this treatment also inhibited the regeneration of a full tail at later stages (**Fig. 1h, i**; analysis details in **Extended Data** Fig. 1b, c and **Methods**). Complementarily, we incubated amputated larvae with the selective nonmuscle myosin II ATPase inhibitor blebbistatin (**Fig. 1j**) to decrease membrane stress and prevent strain stiffening. *i*AFM measurements confirmed that these incubations effectively prevented prospective bud stiffening, and morphological analyses revealed clear impairment of tail regeneration (**Fig. 1k–m**). Together, these results demonstrate that strain stiffening of prospective bud cells during the transition from wound healing to bud formation is required for tail regeneration.

To gain further mechanistic insights into the contribution of prospective bud stiffening to state transitions during regeneration, we next studied how prospective bud cells sense tissue stiffening. Stretch-activated channels have been linked to mechanosensing in epithelial tissues^33–35^. In line with this, our gene expression analysis indicated that the ion channel *piezo1* is highly expressed during the formation of a prospective bud, unlike *piezo2* or members of other ion channel families (i.e., *trpv* and *trpc*), whose expression is lower (**Fig. 2a; Suppl Fig. 3a, b)**. Consequently, we focused on the potential requirement of Piezo1 for regeneration. To this end, we first validated the localization of the endogenous Piezo1 protein to the membrane of prospective bud cells via immunofluorescence (**Fig. 2b**). The functional requirement of Piezo1 for tail regeneration was subsequently assessed via targeted microinjections of a widely validated antisense oligo that induces Piezo1 knockdown in *Xenopus* (Piezo1-MO; **Fig. 2c**)^26,36,37^. Piezo1 loss of function impaired the regeneration of a full tail (**Fig. 2d, e**), indicating its requirement for this process. As a complementary approach and to gain temporal insights into the role of Piezo1 in the transition across cell states during regeneration, we incubated amputated larvae with GsMTx4 (**Fig. 2f**), a peptide that selectively inhibits Piezo1 and TRPs activity in *Xenopus*^26,36,38,39^. Amputated larvae incubated with GsMTx4 during the transition from wound healing to regenerative bud formation strongly inhibited tail regeneration (**Fig. 2g, h, j**). However, this effect was not observed in larvae incubated with GsMTx4 after the regenerative bud had already formed (**Fig. 2g, i, j**). Together, these results show that Piezo1 mechanosensing is required for regeneration during the transition from wound healing to regenerative bud formation.

**Fig. 2.**
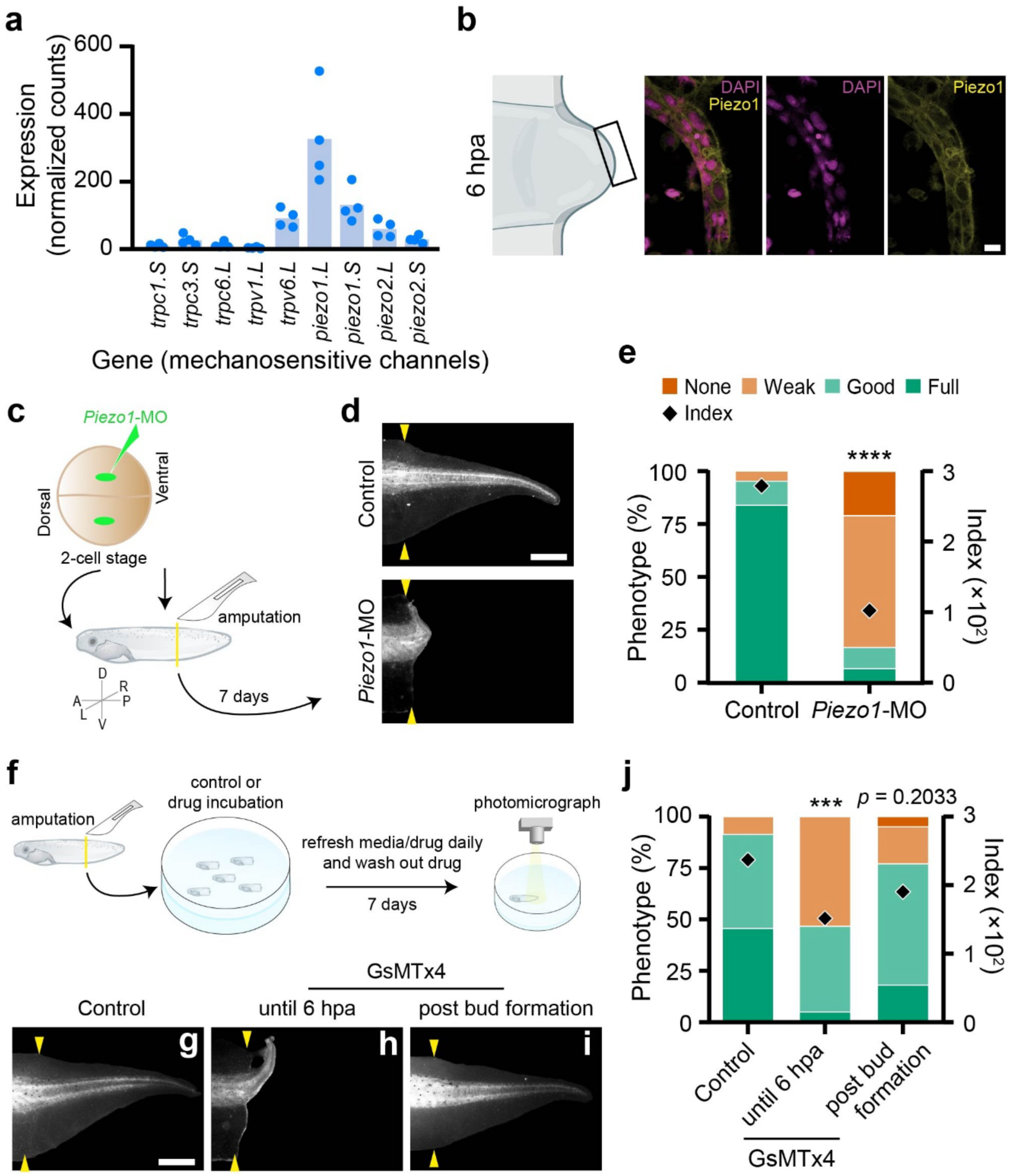
Piezo1 is required for regeneration at the transition from wound healing to regenerative bud formation. **a**, List of stretch-activated ion channels unveiled by RNA-seq of isolated prospective regenerative bud (1 and 6 hpa) and evaluated by DESeq2. *Piezo1* is highly expressed. **b**, Piezo1 membrane localization. Left panel, scheme of the prospective regenerative bud with imaged area (black square); Right panels, confocal slices of immunofluorescence against Pioezo1, with the nuclei counterstained with DAPI. Scale bar, 30 µm. **c–e**, *Piezo1* morpholino (*Piezo1*-MO) inhibits regeneration. **c**, Scheme of morpholino microinjection at 2-cell stage embryos and amputation at stage NF41. A, anterior, P, posterior; D, dorsal; V, ventral; L, left; R, right. **d**, Tails at 7 dpa. Yellow arrowheads, amputation plane. Scale bar, 500 µm. **e**, Quantification of regeneration efficiency. Two-tailed Fisher’s exact test, \*\*\*\**p* < 0.0001, *n*_Control_ = 36, *n_Piezo1_*_-MO_ = 28 tails. **f–j**, Piezo1 inhibitor GsMTx4 impairs regeneration at the transition. **f**, Scheme of the regeneration assay. **g–i**, Tails at 7 dpa. Incubation with 12 µM GsMTx4 until 6 hpa or from 24 to 48 hpa (post bud formation condition). Scale bar, 500 µm. **c**, Quantification of regeneration efficiency. Two-tailed Fisher’s exact test, \*\*\**p* = 0.0004, *p* = 0.2033, *n*_Control_ = 34, *n*_GsMTx4 until 6 hpa_ = 18, *n*_GsMTx4 post bud formation_ = 17 tails. **b,d,g–i**, Representative examples from at least three independent experiments; CI = 95%.

Next, we explored how prospective bud cells translate mechanical signals into a competent regenerative program. Previous work has indicated that Yap1 (Yes-associated protein 1), a transcriptional coactivator of the Hippo pathway, which translates mechanical stress into a transcriptional response^40–43^, is required for *Xenopus* limb and tail regeneration^44,45^. Besides these observations, whether mechanical signals activate Yap1 in these regenerative contexts remains elusive^46^. One of the readouts of Yap1 activity is its nuclear translocation^41,43^. Accordingly, our endogenous Yap1 immunofluorescence analyses revealed a clear increase in Yap1 nuclear localization in prospective bud cells from 1 to 6 hpa (**Fig. 3a, b**). Importantly, this Yap1 translocation coincided with the time at which prospective bud tissues experienced a steeper increase in stiffness (**Fig. 1d, e**), suggesting mechanical control. Indeed, inhibition of prospective bud stiffening prevented Yap1 nuclear translocation (**Fig. 3c, d**), confirming that tissue mechanics influence Yap1 nuclear localization in prospective bud cells. Then, to dissect the temporal requirement of Yap1 for cell state transitions during regeneration, we incubated amputated larvae with verteporfin (**Fig. 3e**), a small molecule that retains Yap1 in the cytosol^47^. We first confirmed the effectiveness of verteporfin by immunofluorescence against Yap1 (**Suppl Fig. 4a, b**). We observed that incubating amputated larvae before bud formation clearly impaired regeneration; however, this effect was not observed when the larvae were incubated after the regenerative bud had already formed (**Fig. 3f–i**). These results indicate that nuclear Yap1 is required for regeneration because it allows wounded tissues to transit into regenerative bud states.

**Fig. 3.**
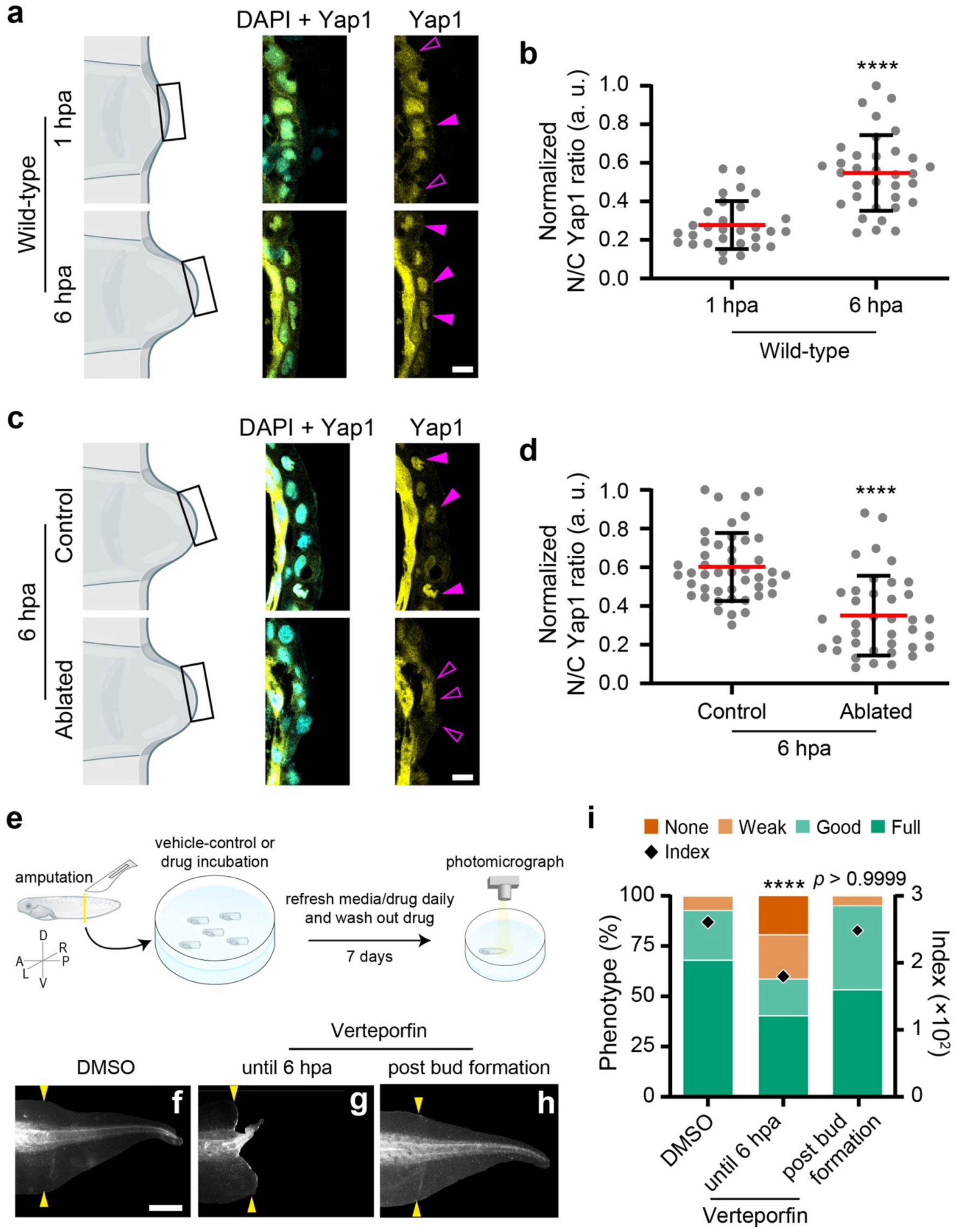
**Yap1 activity is mechanically controlled and is required for the transition from wound healing to regenerative bud formation. a,b**, Yap1 nuclear localization is present and builds up at the transition. **a**, Left panels, schemes of the prospective regenerative bud with imaged area (black squares); Right panels, confocal slices of immunofluorescence against Yap1, with the nuclei counterstained with DAPI. Filled magenta arrowheads, high nuclear localization of Yap1; outlined magenta arrowheads, low nuclear localization of Yap1. Scale bar, 30 µm. **b**, Quantification of Yap1 nucleo-cytoplasmic (N/C) ratio. Red lines represent mean and error bars standard deviation. Two-tailed Student’s *t*-test with Welch’s correction, \*\*\*\**p* < 0.0001, *n*_1 hpa_ = 30, *n*_6 hpa_ = 34 cells. **c,d**, Tissue ablation-mediated lowering of tissue stiffening reduces nuclear localization of Yap1 at the transition. **c**, Left panels, schemes of the prospective regenerative bud with imaged area (black squares); Right panels, confocal slices of immunofluorescence against Yap1, with the nuclei counterstained with DAPI. Scale bar, 30 µm. **d**, Quantification of Yap1 N/C ratio. Red lines represent mean and error bars standard deviation. Two-tailed Student’s *t*-test, \*\*\*\**p* < 0.0001, *n*_Control_ = 45, *n*_Ablated_ = 38 cells. **e–i**, Yap1 inhibitor Verteporfin impairs regeneration at the transition. **e**, Scheme of the regeneration assay. A, anterior, P, posterior; D, dorsal; V, ventral; L, left; R, right. **f–h**, Tails at 7 dpa. Incubation with 2 µM Verteporfin until 6 hpa or from 2 to 7 dpa (post bud formation condition). Yellow arrowheads, amputation plane. Scale bar, 500 µm. **c**, Quantification of regeneration efficiency. Two-tailed Fisher’s exact test, \*\*\*\**p* < 0.0001, *p* > 0.9999, *n*_Control_ = 65, *n*_Verteporfin until 6 hpa_ = 80, *n*_Verteporfin post bud formation_ = 37 tails. **a,c,f–h**, Representative examples from at least three independent experiments; CI = 95%.

Yap1 inhibition (**Fig. 3e–i**) recapitulates the temporal requirement observed upon Piezo1 inhibition (**Fig. 2f–j**). However, whether these molecules operate in parallel or as part of a hierarchical cascade remains to be addressed. Since Piezo1 senses mechanical stress at the cell membrane and Yap1 transduces mechanical stimuli from the cytosol into the nucleus, we next analysed whether Yap1 acts downstream of Piezo1. To explore this, we assessed the impact of Piezo1 inhibition on Yap1 subcellular localization by incubating amputated larvae with GsMTx4. We observed that Piezo1 inhibition prevented both Yap1 nuclear translocation (**Fig. 4a, b**) and tail regeneration (**Fig. 4c, d**). However, coincubation of GsMTx4-treated larvae with PY-60^48^, a chemical activator of Yap1 (**Suppl Fig. 4c, d**), led to both Yap1 nuclear localization and consistent rescue of tail regeneration (**Fig. 4a–d**). In contrast, the impact of Yap1 inhibition on regeneration was not rescued when verteporfin-treated larvae were coincubated with Yoda1^37,49,50^, an activator of Piezo1 (**Fig. 4e, f**). Together, and without ruling out parallel effects, these results indicate that Yap1 works downstream of Piezo1 by eventually relaying tissue stiffening into a transcriptional response that allows wounded epithelia to transit into regenerative states.

**Fig. 4.**
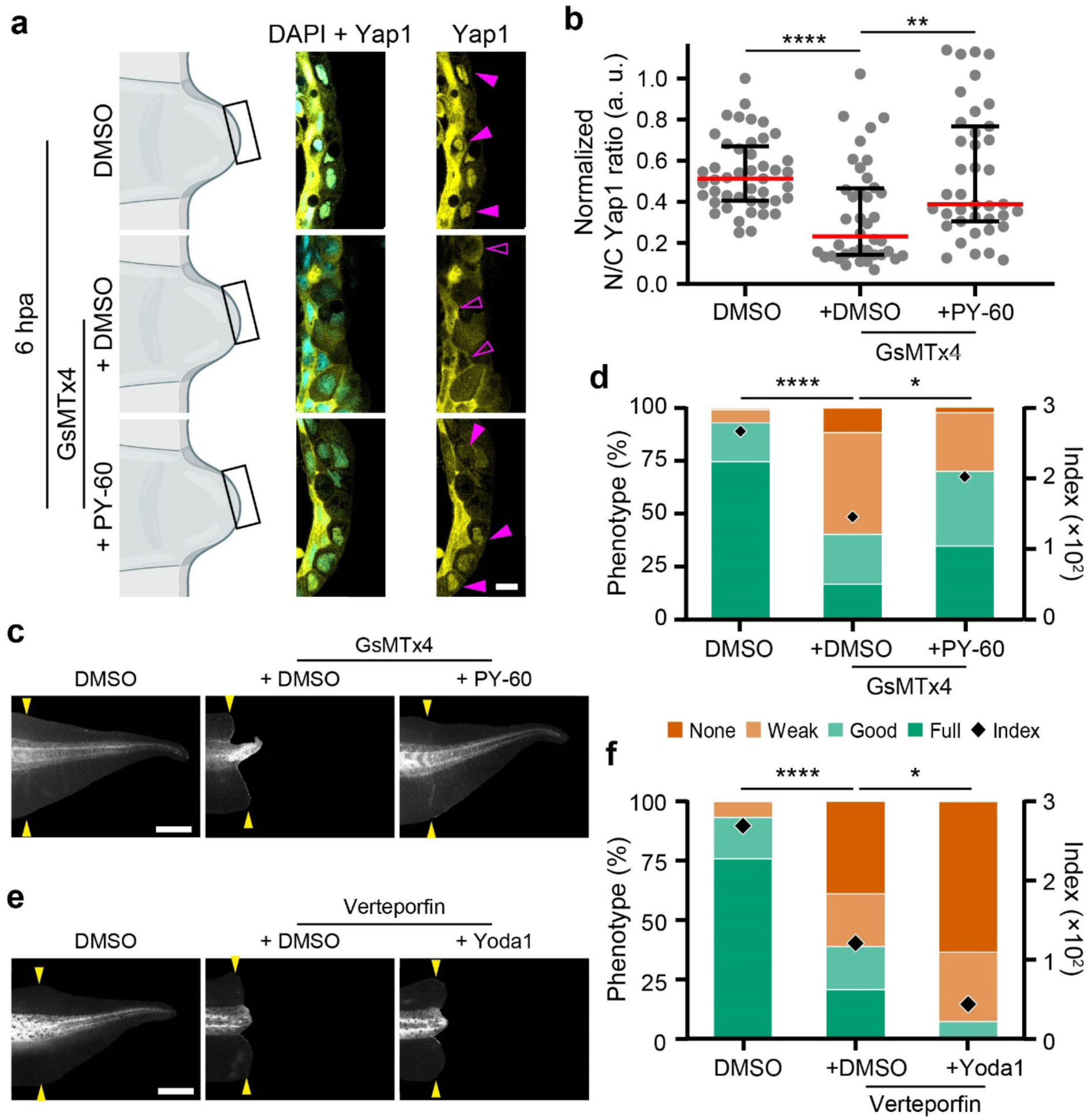
**Yap1 acts downstream of Piezo1 to modulate regeneration. a,b**, Piezo1 modulates Yap1 subcellular localization. **a**, Left panels, schemes of the prospective regenerative bud with imaged area (black squares); Right panels, confocal slices of immunofluorescence against Yap1, with the nuclei counterstained with DAPI. Incubation with 0.1% DMSO, 10 µM GsMTx4 and 1 µM PY-60. Filled magenta arrowheads, high nuclear localization of Yap1; outlined magenta arrowheads, low nuclear localization of Yap1. Scale bar, 30 µm. **b**, Quantification of Yap1 nucleo-cytoplasmic (N/C) ratio. Red lines represent median and error bars interquartile range. Kruskal-Wallis test with Dunn’s multiple comparisons test, \*\**p* = 0.0072, **** *p* < 0.0001, *n*_DMSO_ = 45, *n*_GsMTx4+DMSO_ = 40, *n* _GsMTx4+PY-60_ = 39 cells. **c,d**, Yap1 activator PY-60 rescues regeneration following the inhibition of Piezo1 with GsMTx4. **c**, Tails at 7 dpa. Incubation with 0.1% DMSO, 10–12 µM GsMTx4 and 1 µM PY-60 up to 24 hpa, then washed out. Yellow arrowheads, amputation plane. Scale bar, 500 µm. **d**, Quantification of regeneration efficiency. Two-tailed Fisher’s exact test, \*\*\*\**p* < 0.0001, \**p* = 0.0125, *n*_DMSO_ = 70, *n*_GsMTx4+DMSO_ = 53, *n* _GsMTx4+PY-60_ = 42 tails. **e,f**, Piezo1 activator Yoda1 does not rescue regeneration following the inhibition of Yap1 with Verteporfin. **a**, Tails at 7 dpa. Incubation with 0.21% DMSO, 2 µM Verteporfin and 10 µM Yoda1 until 24 hpa, then washed out. Yellow arrowheads, amputation plane. Scale bar, 500 µm. **b**, Quantification regeneration efficiency. Two-tailed Fisher’s exact test, \*\*\*\**p* < 0.0001 and \**p* = 0.0134, *n*_DMSO_ = 33, *n*_Verteporfin+DMSO_ = 22, *n*_Verteporfin+Yoda1_ = 26 tails. **a,c,e**, Representative examples from at least three independent experiments; CI = 95%.

To test this idea, we performed RNA sequencing of control and verteporfin-treated prospective regenerative buds isolated at 6 hpa, the stage at which we observed strain stiffening to reach a maximum (**Suppl Fig. 5a;** details in **Methods**). Differential gene expression analyses revealed that several genes that are known targets of Yap1^41,42,51,52^ were affected after Yap1 nuclear translocation was prevented (**Suppl Fig. 5b, c**), confirming the quality of our libraries. Further analyses revealed a number of transcripts that encode proteins involved in signalling pathways (e.g., FGF, Notch, Wnt, and BMP) and biological processes (e.g., the cell cycle, migration, fate decision, and metabolism), which are known to be essential for regeneration^3,5–7,21,53,54^ (**Fig. 5a–c**). These results indicate that Yap1 nuclear activity is required to support the acquisition of a gene expression profile that allows prospective bud cell transition into a competent regenerative state.

**Fig. 5.**
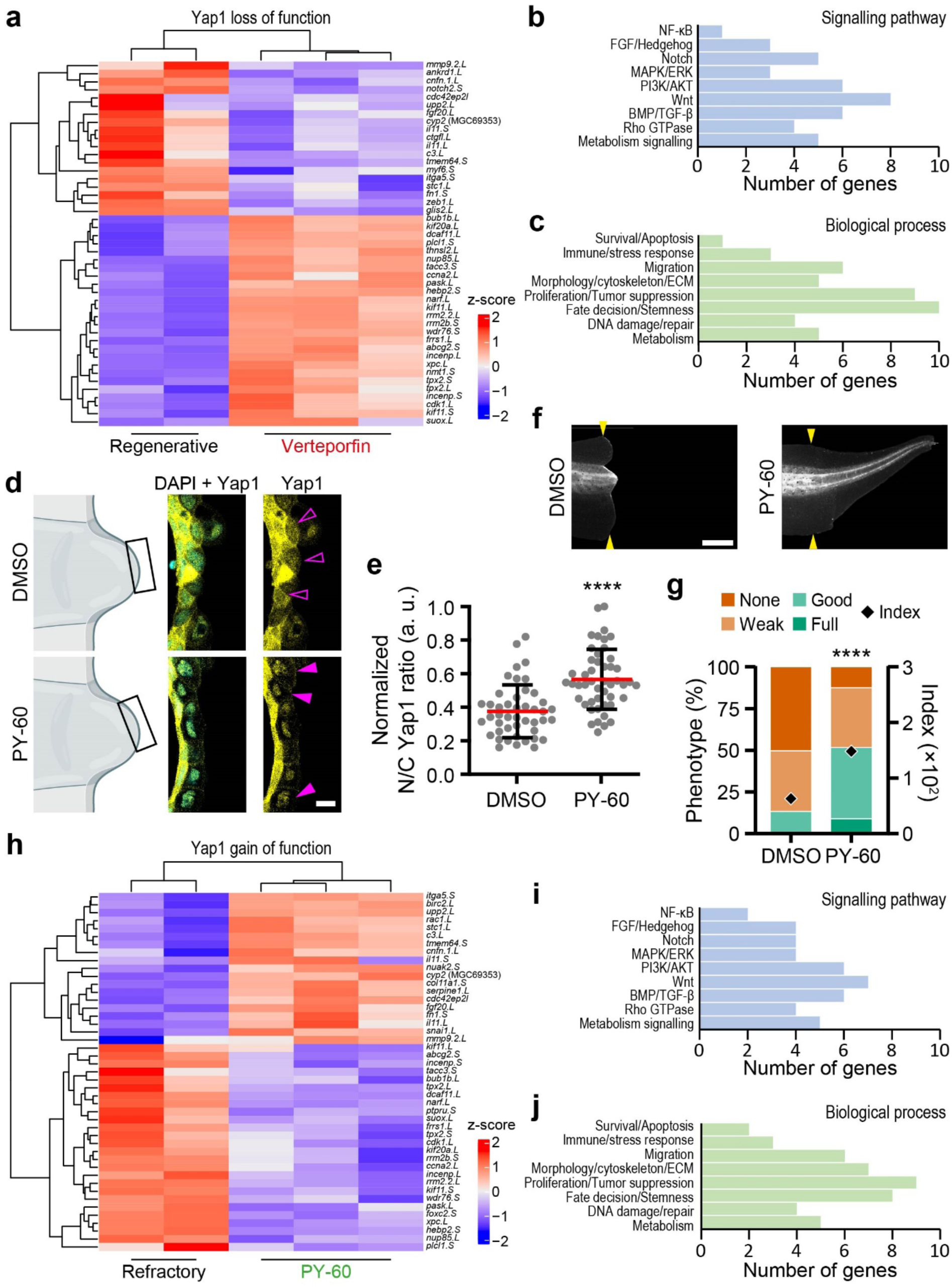
Yap1 is required and sufficient to relay strain stiffening into regenerative competence. **a–c**, Yap1 transcriptional program includes genes (**a**), signalling pathways (**b**) and biological processes (**c**) known to be important for regeneration. **a**, Heatmap with expression levels of Yap1 target genes following Yap1 inhibition (regenerative wild-type vs. 2 µM Verteporfin for 6 hpa). Each column is a different sample. Positive (red) values mean upregulated gene and negative (blue) values mean downregulated gene. Histograms of Yap1 target genes (from **a**) clustered by signalling pathways (**b**) or biological processes (**c**). **d,e**, Yap1 activator PY-60 augments nuclear localization of Yap1 in tails from refractory period (6 hpa). **d**, Left panels, schemes of the prospective regenerative bud with imaged area (black squares); Right panels, confocal slices of immunofluorescence against Yap1, with the nuclei counterstained with DAPI. Incubation with 0.1% DMSO and 1 µM PY-60. Filled magenta arrowheads, high nuclear localization of Yap1; outlined magenta arrowheads, low nuclear localization of Yap1. Scale bar, 30 µm. **e**, Quantification of Yap1 nucleo-cytoplasmic (N/C) ratio. Red lines represent mean and error bars standard deviation. Two-tailed Student’s *t*-test, \*\*\*\**p* < 0.0001, *n*_Refractory DMSO_ = 45, *n*_Refractory PY-60_ = 48 cells. **f,g**, Yap1 activator PY-60 induces regeneration in the otherwise nonregenerative tails of refractory period. **f**, Tails at 7 dpa. Incubation with 0.1% DMSO or 1 µM PY-60 until 24 hpa. Yellow arrowheads, amputation plane. Scale bar, 500 µm. **g**, Quantification of regeneration efficiency. Two-tailed Fisher’s exact test, \*\*\*\**p* < 0.0001, *n*_Refractory DMSO_ = 56, *n*_Refractory PY-60_ = 48 tails. **h–j**, Yap1 activation in the refractory period gains a regenerative transcriptional program in otherwise nonregenerative tails. **h**, Heatmap with expression levels of Yap1 target genes following Yap1 activation (refractory wild-type vs. 1 µM PY-60 for 6 hpa). Histograms of Yap1 target genes (from **h**) clustered by signalling pathways (**b**) or biological processes (**c**). **d,f**, Representative examples from at least three independent experiments; CI = 95%.

In this context, we next asked whether the activation of this mechanosensitive cascade is sufficient to bring wounded tissues into a regenerative-like state in nonregenerative contexts. For this purpose, we took advantage of the *Xenopus laevis* regeneration incompetent period (also known as the refractory period), – a later developmental stage (NF45–47) in which larvae fail to regenerate after amputation (**Suppl Fig. 6a, b**). We first analysed the Yap1 distribution in the wild-type refractory period as a readout of mechanosensitive activation. Immunofluorescence analyses revealed that Yap1 localization was not biased towards the nuclei of prospective bud cells, in contrast to the clear nuclear Yap1 signal detected at the regenerative stage (NF41) (**Suppl Fig. 6c, d**). Thus, to assess whether Yap1 nuclear localization is sufficient to drive regeneration, we incubated refractory period larvae with the Yap1 activator PY-60. This treatment led to a clear translocation of Yap1 into the nuclei of wounded refractory cells, which are topologically related to tissues in which a prospective bud would form in regenerative stages (**Fig. 5d, e**). Notably, Yap1 activation also strongly increased regenerative competence in these otherwise nonregenerative tissues (**Fig. 5f, g**). Indeed, RNA sequencing of “prospective regenerative buds” isolated from the refractory period (**Suppl Fig. 7a**) confirmed that the observed gain of regeneration was accompanied by the induction of Yap1 target genes (**Suppl Fig. 7b, c**). In addition, we observed the activation of a regenerative gene expression profile (**Fig. 5h–j**). These results complement those obtained with verteporfin (**Fig. 5a–c**), as markers downregulated by Yap1 inhibition are upregulated by PY60-mediated activation (**Suppl Fig. 7d**). Taken together, these results suggest that Yap1 activation confers regenerative incompetent tissues with genetic profiles that support the reactivation of pathways required for wounded tissues to transition into bud-like states, resulting in the acquisition of regenerative competence.

### Outlook

The stages through which regenerating tissues transit during epimorphic regeneration are fairly well studied; however, how the transitions across these phases are synchronized remains comparatively less understood. Addressing this fundamental aspect of regeneration is relevant, as it may hold the key to reactivating regenerative programs in nonregenerative tissues, which is a major challenge in the field. Our work contributes to this goal, as we reveal an otherwise “hidden” stage of regeneration, which involves the transition from wound healing to regenerative bud formation. This transient stage is marked by a steep mechanical build-up in the prospective bud, an event required for these cells to acquire their characteristic regenerative competence (**Fig. 6a**). The formation of a competent regenerative bud represents a critical transition in regeneration, as this transition occurs when and where tissues acquire their regenerative capacity^3,5,21^. Indeed, these are the stages and tissues in which the recently discovered Regeneration Organising Cells (ROCs) and Regeneration Initiating Cells (RICs) are described to operate^23,24^. Hence, the temporal and positional relevance of our findings and their potential to provide mechanistic insights into how tissues acquire regenerative competence.

**Fig. 6.**
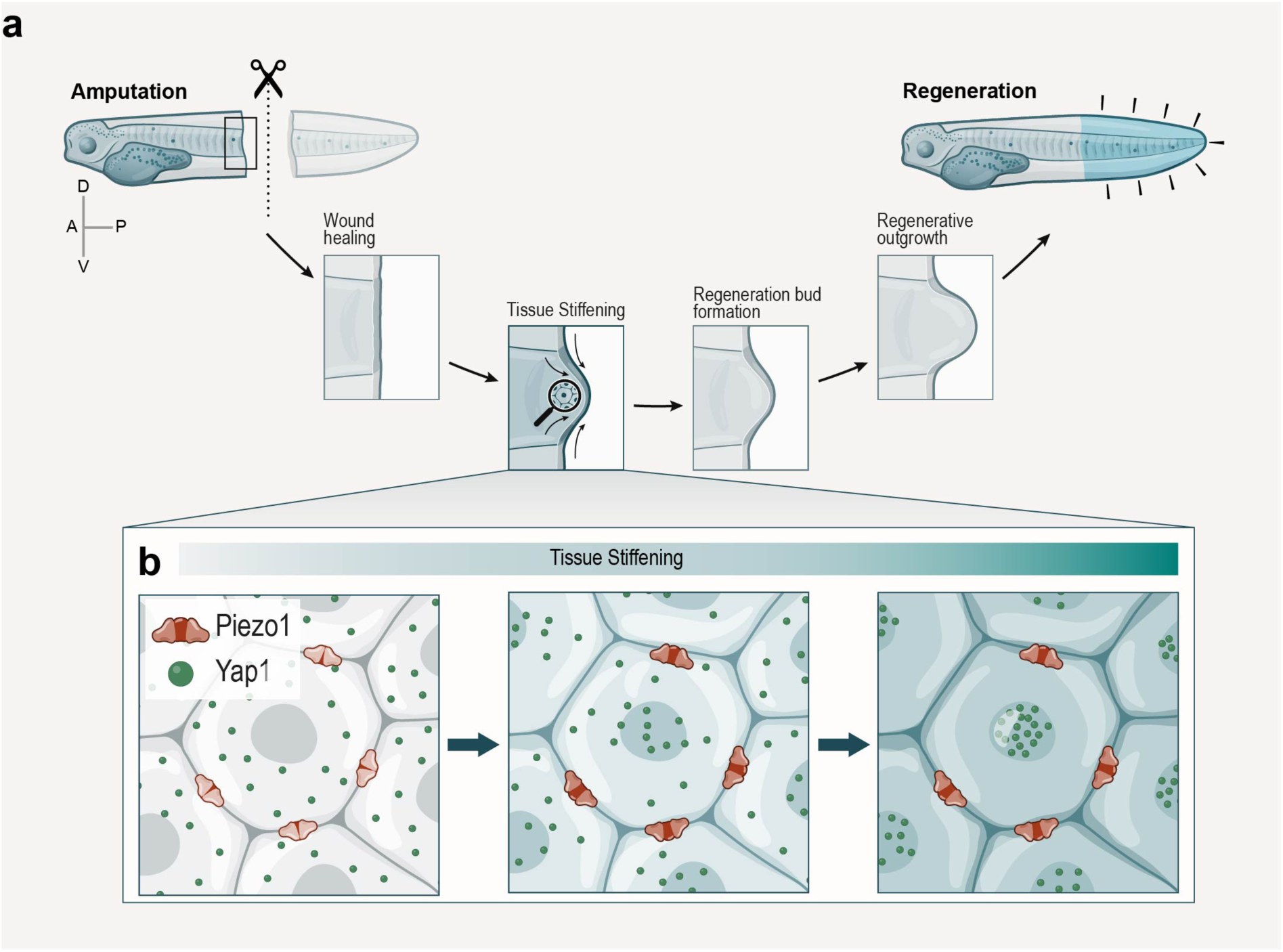
Summary model: Mechanomolecular control of cell state transitions during regeneration via Piezo1–driven Yap1 activation. **a**, Schematic of the tail regeneration phases highlighting a newly identified state transition– prospective bud tissue strain stiffening from wound healing to regenerative bud formation. **b**, Zoom-in of the prospective regenerative bud at the wound healing to bud transition, highlighting Piezo1–Yap1 mechanosensitive axis. In this transition, the tissue strain stiffening leads to a Piezo1-dependent nuclear localization of Yap1. This mechanomolecular axis leads to the transcription of a regenerative program that allow wounded epithelial tissues to transit into a competent bud state to drive epimorphic regeneration.

Moreover, we found that prospective bud stiffening triggers a mechanosensitive cascade in which Piezo1 drives the activation of Yap1. This mechanosensitive cascade drives prospective bud stiffening into a genetic response, which is not only necessary but also sufficient to confer the regenerative capacity of prospective bud cells (**Fig. 6b**). While interesting *per se*, the mechanism by which Piezo1 influences Yap1 nuclear localization remains to be addressed. Yap1 is translocated from the cytosol into the nuclei of cells in response to mechanical stress. The interaction of Yap1 with 14-3-3 proteins is part of the mechanism by which Yap1 is retained in the cytosol^43,47,55^. Considering that Piezo1 transduces mechanical stress via calcium influx and that some studies related calcium to 14-3-3 protein activity^56^, an interesting possibility is that Piezo1 controls Yap1 activation via 14-3-3 regulation. Thus, our work opens new research avenues aimed not only at revealing how Piezo1 controls Yap1 activity but also at exploring whether and how this Piezo1-Yap1 mechanosensitive axis operates in other biological contexts.

Importantly, we observed that this new Piezo1-Yap1 mechanosensitive axis is not only required but also sufficient to support a transcriptional program that drives regeneration. We observed that Yap1 is inactive at nonregenerative stages and that its chemical activation strongly induces regeneration. While this is a unique observation, the mechanism by which Yap1 is retained outside of the cell nucleus during refractory stages remains unknown. One possibility is that Piezo1 is not expressed in those stages; however, our sequencing data rule out that possibility. Another idea is that the signalling cascade that works downstream of Piezo1 to activate Yap1 may not be present or functional in these stages (i.e., metabolic states or other related signalling pathways^57,58^) or that tissue mechanics do not develop at these stages. In line with this, a recent report showed that, by turning on an evolutionarily disabled genetic switch, regeneration of mammalian ears can be reactivated^59^. Additionally, the jaws of mice require mechanical stimulation to reactivate developmental programs and regeneration^60^. Thus, another possibility is that tissue mechanics may influence the activity of that switch. While further experimentation will dissect this point, our observations emphasize the idea that studying how tissues transit from one regenerative state to another can provide insights into how to drive *de novo* regeneration in nonregenerative contexts.

Broadly, our discoveries highlight the relevance of integrating endogenous tissue mechanics with molecular frameworks to actuate morphogenesis by synchronizing cell state transitions.

## Author contributions

E.H.B. conceptualized the project with inputs from F.F. F.F. and E.H.B. developed the research.

F.F. performed most of the experiments and data analyses with some help from Q.T., I.F. and E.H.B. J.A.E. performed the AFM experiments with help from F.F. and E.H.B. A.G.K. performed the initial bioinformatic analyses. F.F. and E.H.B. prepared the manuscript and figures with input from all the authors. E.H.B. supervised the project.

## Acknowledgements

We thank Dr. Patricia Ramos for the comments on the manuscript. We acknowledge the services of the IGC’s and TUD’s Imaging, Genomics, and Aquatic animal facilities. Jose Grau from Genomica Australis for bioinformatic services. Julien Marcetteau for help with illustrations. Work at the Barriga laboratory was supported by the European Research Council Starting Grant (ERC-StG) under the European Union’s Horizon 2020 research and innovation programme, Grant agreement No. 950254; the European Molecular Biology Organization (EMBO) Installation Grant, Project No. 4765; the EMBO Young Investigator Programme, Project No. 5248; the La Caixa Junior Leader Incoming, No. 94978; the Fundação Calouste Gulbenkian (FCG), start-up grant I-411133.01; and an EMBO postdoctoral fellowship ALTF 27-2020 (to FF). The Barriga laboratory is also funded by the Deutsche Forschungsgemeinschaft (DFG, German Research Foundation) under Germany’s Excellence Strategy (EXC 2068, 390729961, Cluster of Excellence Physics of Life of TU Dresden).

## Data availability

All data supporting this study is available throughout the manuscript. All other relevant information is available from the corresponding author upon reasonable request.

## Methods

### Ethics statement

Animal experimental procedures and euthanasia were approved by the Ethics Committee and Animal Welfare Body (ORBEA) of the Instituto Gulbenkian de Ciência (IGC). Procedures complied with the legislations from Portugal (Decreto-Lei n° 113/2013) and from the European Union (Directive 2010/63/EU) for animal experimentation and welfare.

### Frog manipulation to obtain tadpoles

The embryos of *Xenopus laevis* (Daudin, 1802) were obtained by *in vitro* fertilization^61^ by injecting human chorionic gonadotropin (Chorulon) to induce ovulation in adult females. Freshly collected oocytes were fertilized by mixing with a sperm solution. Embryos were maintained at 12 and 23 °C and staged according to the normal table of development^62^. Amputation assays were performed at tadpole stages: NF40–45.

### Tail regeneration assay

Tadpoles were anesthetized in 1 mM tricaine methanesulfonate (MS222; Pharmaq, 1004671) at stages NF40–41 (regenerative) or stage NF45 (refractory period) and amputated under a dissecting microscope as previously described^22,63^. After washing out the anaesthetic, animals were transferred into the specified experimental condition. Drug incubations were refreshed daily, and pictures of the tails were taken at 7 days postamputation (dpa) (**Suppl Fig. 1a**). To quantify the regenerative ability of the tadpoles under any treatment four predefined phenotypic categories were stablished considering axial outgrowth and patterning^22,63^: *Full*, complete tail regeneration that is identical to uncut tadpoles; *Good*, robust tail regeneration but with minor defects, such as missing fin or curved body axis; *Weak*, poor tail regeneration with abnormal or hypomorphic regenerates; *None*, absent tail regeneration (**Suppl Fig. 1b**). Then the frequencies (*f*) of each category within the population were used to calculate a regeneration index as follow^22,23,63^ (**Suppl Fig. 1c**):

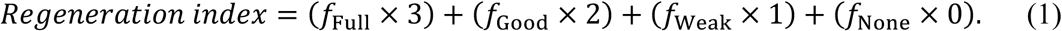

This index ranges from 0 to 300, where the extremes 0 means absent regeneration and 300 means complete regeneration. For guidance, an averaged index of > 250 represents nearly unimpaired regeneration, and an averaged index of < 150 represents highly impaired regeneration.

In addition, images were taken at 72 hpa to see the effect on regeneration^64^ by measuring the area of regenerated tail normalized to the intact somite area. These measurements were performed using Fiji software v1.54p (https://imagej.net/software/fiji/).

### Microinjection and pharmacologic modulations

mRNA and morpholino microinjections in both blastomeres at 2-cells stage. Glass needles were calibrated to deliver 10 nl on a gas pulse of 20 psi for 0.2 s. 250 pg of nuclear RFP and/or membrane GFP, or 34 ng of *Piezo1* morpholino (CACAGAGGACTTGCAGTTCCATCCC)^26^ was injected per blastomere. RNAs were obtained by in vitro transcription with the mMESSAGE mMACHINE SP6 kit (Thermo-Fisher, AM1340) while morpholinos were obtained as previously described^36^.

GsMTx4^65^ (Smartox Biotechnology, 08GSM001) was used at 12 µM unless otherwise specified, blebbistatin (Abcam, AB120425) was used at 50 µM, verteporfin^47^ (Sigma-Aldrich, SML0534) was used at 2 µM and PY-60^48^ (MedChemExpress, HY-141644) was used at 1 µM. GsMTx4 was stocked with water, verteporfin and PY-60 were stocked with dimethyl sulfoxide (DMSO) and stored at −80 °C. Blebbistatin was stocked in DMSO and stored at −20 °C. Prior to use, drugs were freshly reconstituted in 0.1× MMR.

### *In vivo* atomic force microscopy measurement

The apparent elastic moduli (referred as stiffness) of tissues were measured using an *in vivo* atomic force microscope (*i*AFM) (**Fig. 1b,c**), as previously detailed^36,66^. Briefly, measurements were performed with the automated Flex-ANA (Nanosurf) AFM device mounted with a *xy* motorized stage, and controlled by an ANA software v1.3 (Nanosurf). We used colloidal AFM cantilevers with a 10 ± 1 µm diameter sphere tip coated with reflective chromium/gold (sQube, CP-qp-SCONT-BSG-B). The cantilevers were mounted onto the AFM device, and, after calibration, their spring constants were calculated with the thermal noise method^67^, ranging from 0.01 to 0.02 N m^−1^. For the measurements, the tadpoles were anesthetized with tricaine 1 mM and immobilized in a custom-made measuring dish filled with 0.1× MMR supplemented or not with vehicle-control or drug. For data acquisition, we used a 10 × 10 µm grid comprised of 5 lines and 5 rows, with the following modulation parameters: maximum indentation force, 10 nN; approach speed, 5 μm s^−1^; retraction speed, 50 μm s^−1^; and sample rate, 2,400 Hz.

### *In vivo* atomic force microscopy analysis

The force–distance curves acquired from each grid (**Suppl Fig. 2a,b**) were fitted to a Hertz model for a spherical indenter, as in the following equation^26,68^:

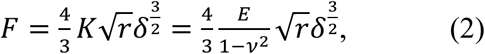

where *F* is the applied force, *E* is the Young’s modulus, *ν* is the Poisson’s ratio, *r* is the indenter radius, *δ* is the indentation depth, and *K* is the apparent elastic moduli (referred in the text as ‘stiffness’ and in the *y* axis of plots as ‘apparent elasticity’). The force–distance curves were selected as previously detailed^66^ (representative example in **Suppl Fig. 2a**). And the apparent elasticity moduli were extracted at maximum indentation depth using AtomicJ software v2.3.1. Then, the median of each embryo was calculated and processed for further statistical analyses.

### Immunofluorescence

Tadpoles were fixed overnight at 4 °C with 3.7% formaldehyde in 0.3% PBS Triton X-100 and, after washing, had their heads excised. The tails were blocked in 10% normal goat serum (NGS) or 2% bovine calf serum (BCS), depending on antibody and incubated at 1 h at room temperature. The primary antibody anti-Piezo1 (Abcam, AB128245) and anti-Yap1 (Abcam, AB56701) were diluted 1:75 in 10% NGS and 1:100 in 2% BCS, respectively. After washing with 0.1% PBS Tween-20, the tails with anti-Piezo1 were incubated for 2 h at room temperature with anti-rabbit Alexa Fluor 555 (Invitrogen, A-21429) at 1:350 in 10% NGS and 1:500 DAPI. The tails with anti-Yap1 had the signal amplified with the tyramide SuperBoost kit (Invitrogen, B40916), following manufacturer’s instructions. Briefly, tails were incubated for 2 h at room temperature with anti-mouse HRP and 1:500 DAPI. After washes, tails were incubated with the Alexa Fluor tyramide reagent for 8 min and then reaction was stopped. Tails from either Piezo1 or Yap1 staining were washed and fixed with 3.7% formaldehyde in PBS for 20 min at room temperature. For tissue clearing, tails were dehydrated in an ascending series of methanol – 25, 50, 75 and 100% – in PBS and left in 100% methanol PBS overnight at 4 °C. Tails were incubated with 1:2 BABB (benzyl alcohol and benzyl benzoate; Sigma-Aldrich, 8.22259 and B6630) for 1 h at room temperature and mounted in BABB for confocal microscopy.

### Microscopy

*Regeneration assay.* Images tadpoles were taken with 0.67× or 2.5× magnifications using an USB Dino-Eye eyepiece camera (Dino-Lite, AM7025X) mounted onto a stereoscope, controlled by DinoCapture v2.0 (Dino-Lite) software. Some images were acquired with a stereoscope (Zeiss Stereo Lumar v12) equipped with an uEye camera (IDS), controlled by Micro-Manager v1.14 (μManager).

*Cell membrane imaging.* A *z* stack (1–2 µm step size) of tadpole tails was acquired with an upright Stellaris 5 (Leica) confocal system, using a HC PL APO 25×/0.95 NA water immersion objective (Leica). The system was controlled by the LAS X (Leica).

*Immunofluorescence.* A *z* stack (1 or 2 µm step size) of tadpole tails was acquired with a Stellaris 5 or inverted LSM 900 (Zeiss) confocal systems, using a HC PL APO 25×/0.95 NA water immersion objective or a Plan-Apochromat 63×/1.40 NA (Zeiss) oil immersion objective with 0.45× optical zoom, respectively. The systems were controlled by LAS X or ZEN Blue edition v3.5 (Zeiss), respectively.

### Tissue ablation

Under a microscope, anesthetized tadpoles were ablated along the dorsal and ventral fins at the intersection with the tail trunk. Ablations were performed from the anterior to posterior axis orientation as in **Fig. 1f**. After ablation the larvae were followed along regeneration and analysed at the indicated stages.

### Microscopy analysis

Yap1 subcellular localization was quantified by the nucleo-cytoplasmic (N/C) ratio as previously described^69^. In brief, we located the confocal slice where the nucleus is larger, as determined by the DAPI channel, and used this to determine the nuclear and cytoplasmatic (surrounding nucleus) ROIs. Then, on the corresponding Yap1 channel, the fluorescence density of the respective ROIs was acquired in Fiji and used to calculate the N/C ratio.

### Image processing

Maximum projections of *z*-stacks were created in Fiji. Other common image adjustments, as contrast and brightness, rotation, pseudocolouring and lookup tables, as well as overlay of text and scales, was performed in Fiji and/or Illustrator 2025 (Adobe). Adjustment of contrast and brightness and the background around the tails of the photomicrographs acquired using the Dino-Eye eyepiece camera were pseudocoloured in black in Photoshop (2025, Adobe), to improve clarity but without interfering with the specimen itself.

### RNA-seq procedures

RNA was extracted from isolated perspective regenerative buds and RNA-seq procedures – per single bud – were performed by the IGC Genomics Unit as follows (**Suppl Fig. 3a**). The extracted RNA quality was assessed in a HS RNA Screen Tape Analysis (Agilent Technologies) and the mRNA libraries were prepared using SMART-Seq2 as previously detailed^70^. Libraries were generated with the Nextera protocol^71^ and their quality assessed by a Fragment Analyzer (AATI). The libraries were sequenced with the NextSeq500 Sequencer (Illumina) using a 75 SE high-throughput kit. Finally, the sequences were extracted in a FastQ format using the bcl2fastq v2.19.1.403 (Illumina).

### RNA-seq analysis

For bioinformatic analysis, the *Xenopus laevis* reference genome (XENLA_10.1) and the corresponding gene annotation (Genome Feature File) were obtained from Xenbase (RRID:SCR_003280). Raw sequencing reads were quality-checked using FastQC v0.12.1 (https://www.bioinformatics.babraham.ac.uk/projects/fastqc/), and filtered using Fastp^72^ v0.23.4 with default parameters. Filtered reads were aligned to the *X. laevis* reference genome using HISAT2^73^ v2.2.1. The alignment files were sorted and indexed using Samtools^74^ v1.21. Gene-level quantification was performed using the featureCounts function from the Subread^75^ package v2.0.6. Ribosomal RNA (rRNA) reads were removed prior to differential expression analysis, which was conducted using the DESeq2^76^ package v4.4 in R (The R Project for Statistical Computing; v4.5.04). Heatmaps were generated using the ComplexHeatmap^77^ package v2.25.1 in R. Plots were visualized using ggplot2 v3.5.2 in R.

### Statistical analysis

No software was used for sample size determination. No blinding was followed in data acquisition and analysis owing to the nature of procedures where unviable and uninjected/misinjected tadpoles were excluded. Tadpoles were randomly allocated to the experimental conditions (annotated in figure captions), and experiments were repeated at least 3 independent times. Outliers were flagged using Grubbs’ test or ROUT method and datasets were tested for normality using the d’Agostino–Pearson, Kolmogorov-Smirnov or Shapiro–Wilk tests. Significances were computed with a Student’s *t*-test (two-tailed *p* value). For multiple comparisons, we used the Brown-Forsythe and Welch ANOVA followed by Dunnett’s multiple comparisons tests (two-tailed *p* value) for normally distributed datasets or Kruskal-Wallis followed by Dunn’s multiple comparisons test (two-tailed *p* value) for datasets without normal distribution. Contingency tables were analysed with the Fisher’s exact test (two-tailed *p* value). Statistical analysis was performed in Prism10 (GraphPad; v10.5.0) with a 95% confidence interval for all analysis and statistical details are annotated in all figure captions.

## EXTENDED DATA FIGURES

**Suppl Fig. 1.**
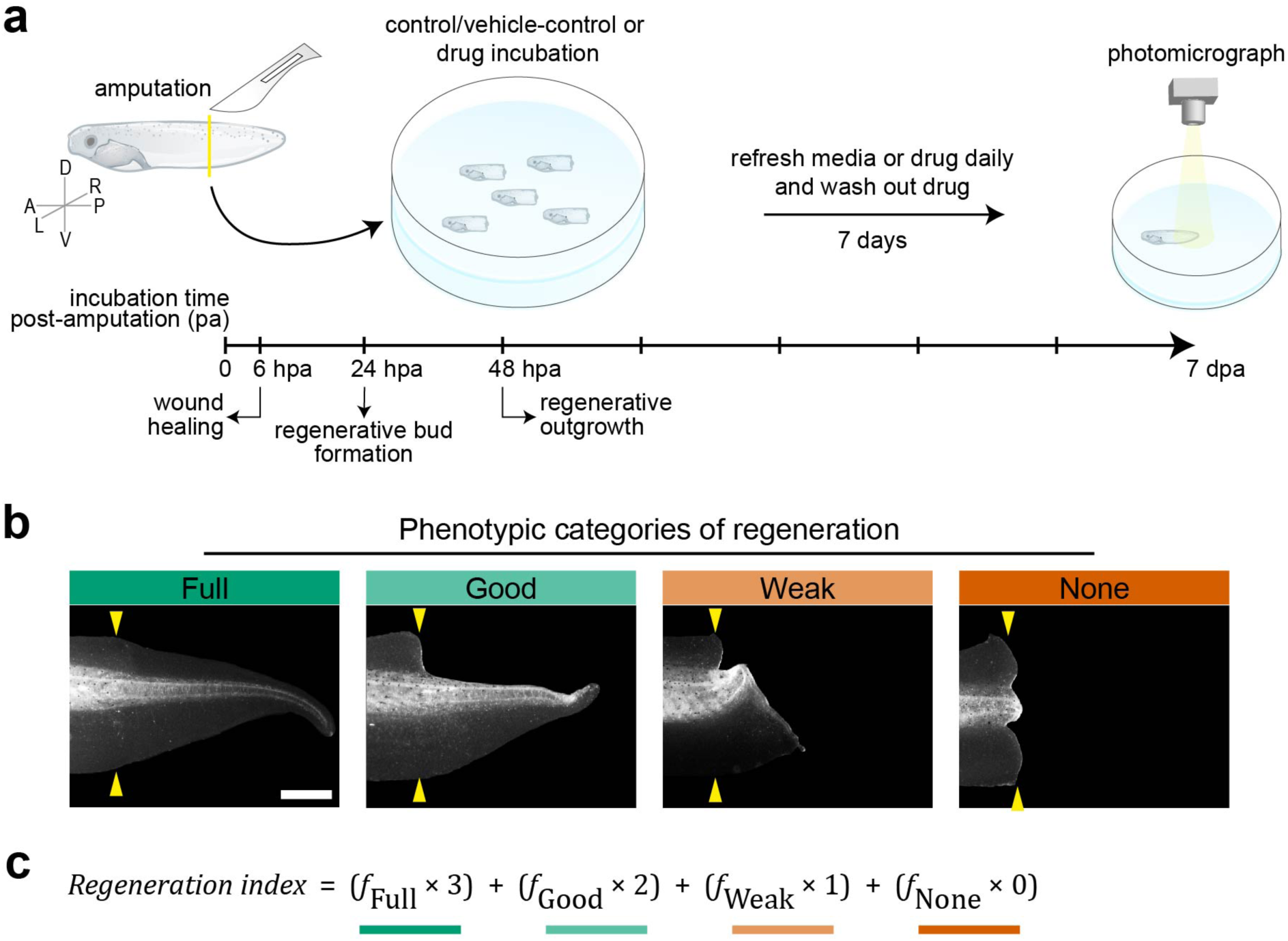
*Xenopus* tadpole tail regeneration assay. **a**, Scheme of the regeneration assay with annotation of the phases of regeneration used for drug incubations. A, anterior, P, posterior; D, dorsal; V, ventral; L, left; R, right. Yellow solid line, amputation plane. **b,c**, Quantitative basis of regeneration efficiency from regeneration assays by means of the regeneration index (defined in **Methods**). **b**, Tails at 7 dpa of each phenotypic category of regeneration used to compute the regeneration index. Scale bar, 500 µm. Representative examples from at least three independent experiments; CI = 95%. **c**, Equation of the regeneration index where *f* is the frequency of regenerates of each phenotypic category. Index ranges from 0, meaning absent regeneration, to 300, meaning complete regeneration.

**Suppl Fig. 2.**
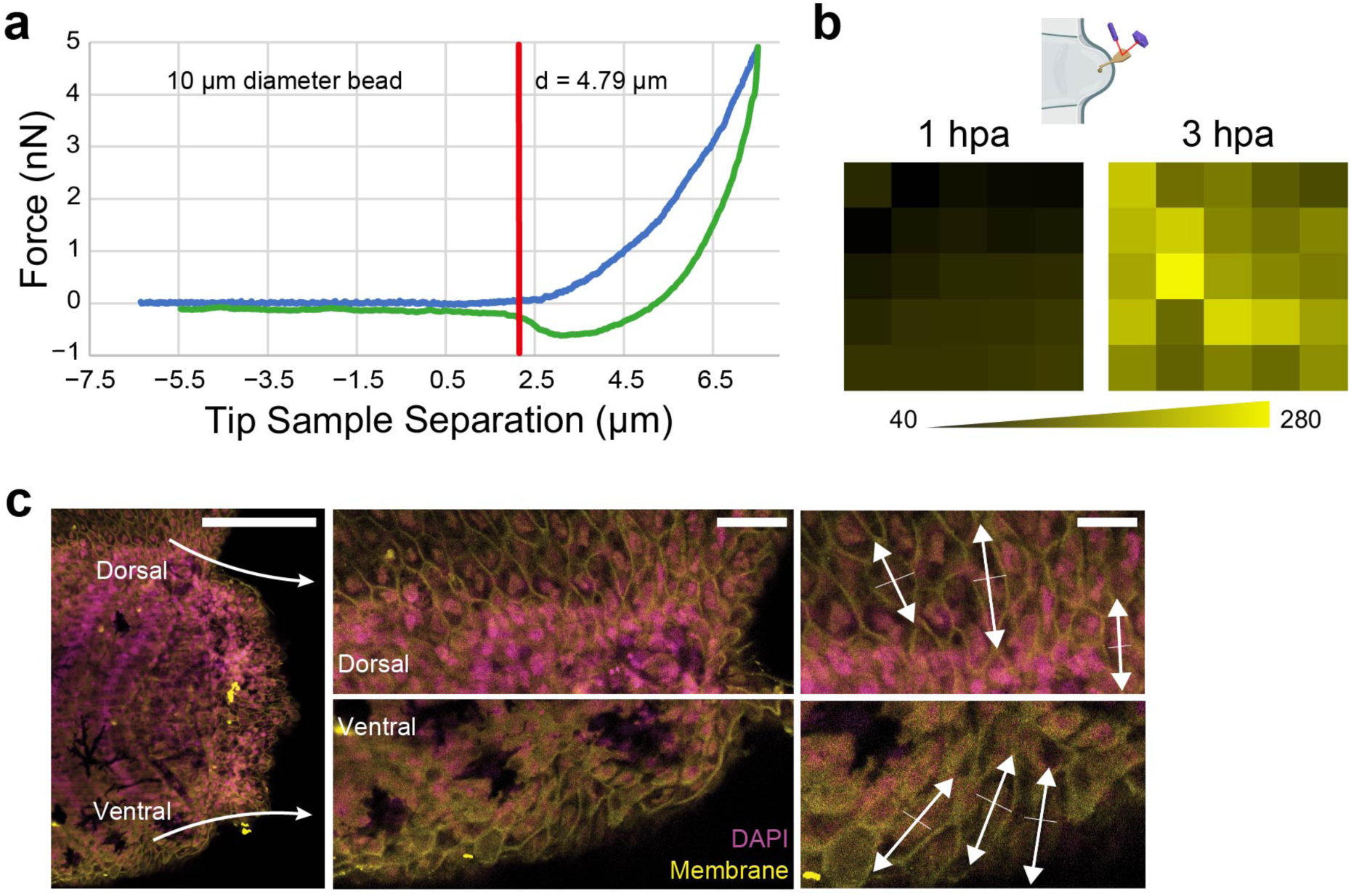
*In vivo* atomic force microscopy measurements and cell membrane stress orientation in the prospective regenerative bud. **a**, Force-distance curve obtained in a typical measurement. **b**, Heatmaps of a 10×10 µm measurement grid acquired every 2 μm, showing the spread of data at 1 and 3 hpa. From these grids, a median data point was calculated and added to the plots presented in the main figures. **c**, Confocal projection showing that the tension is distributed along the dorsoventral axis at the transition from wound healing to regenerative bud formation (6 hpa). Right panel, zoom ins highlighting the dorsoventral cell length (white double-headed arrows) relative to the anteroposterior cell length (white solid lines). Scale bars, 200 µm (left panel), 50 µm (middle panel) and 30 µm (right panel). **a–c**, Representative examples from at least three independent experiments; CI = 95%.

**Suppl Fig. 3.**
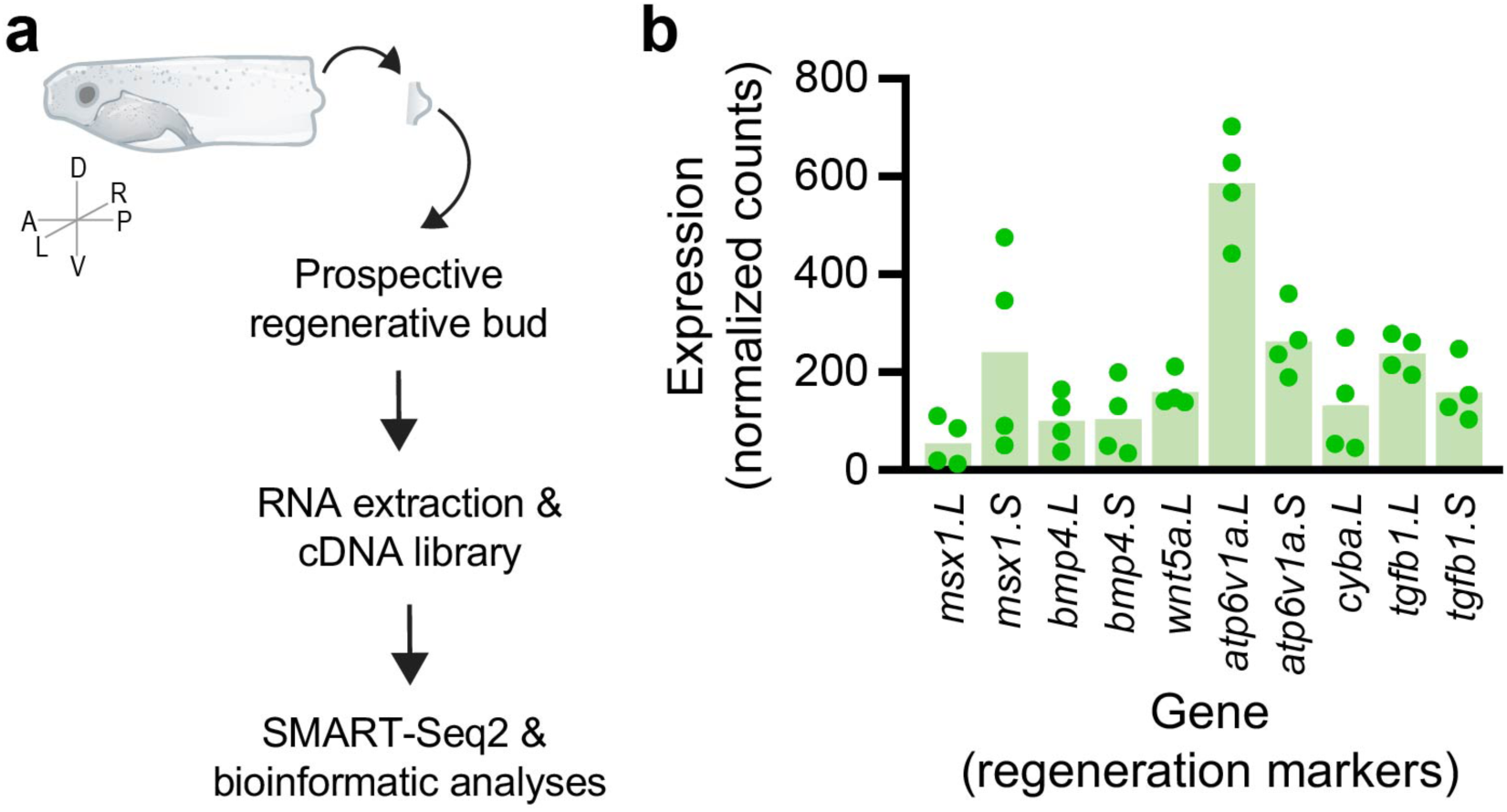
RNA-seq of wild-type isolated prospective regenerative buds at 1 and 6 hpa. **a**, Scheme and flowchart for the collection and processing of isolated prospective regenerative buds (stage NF41) for SMART-Seq2 RNA sequencing. A, anterior, P, posterior; D, dorsal; V, ventral; L, left; R, right. **b**, List of marker regeneration genes at 1 and 6 hpa, evaluated by DESeq2. Majority of regeneration markers are enriched, validating the quality of our RNA libraries.

**Suppl Fig. 4.**
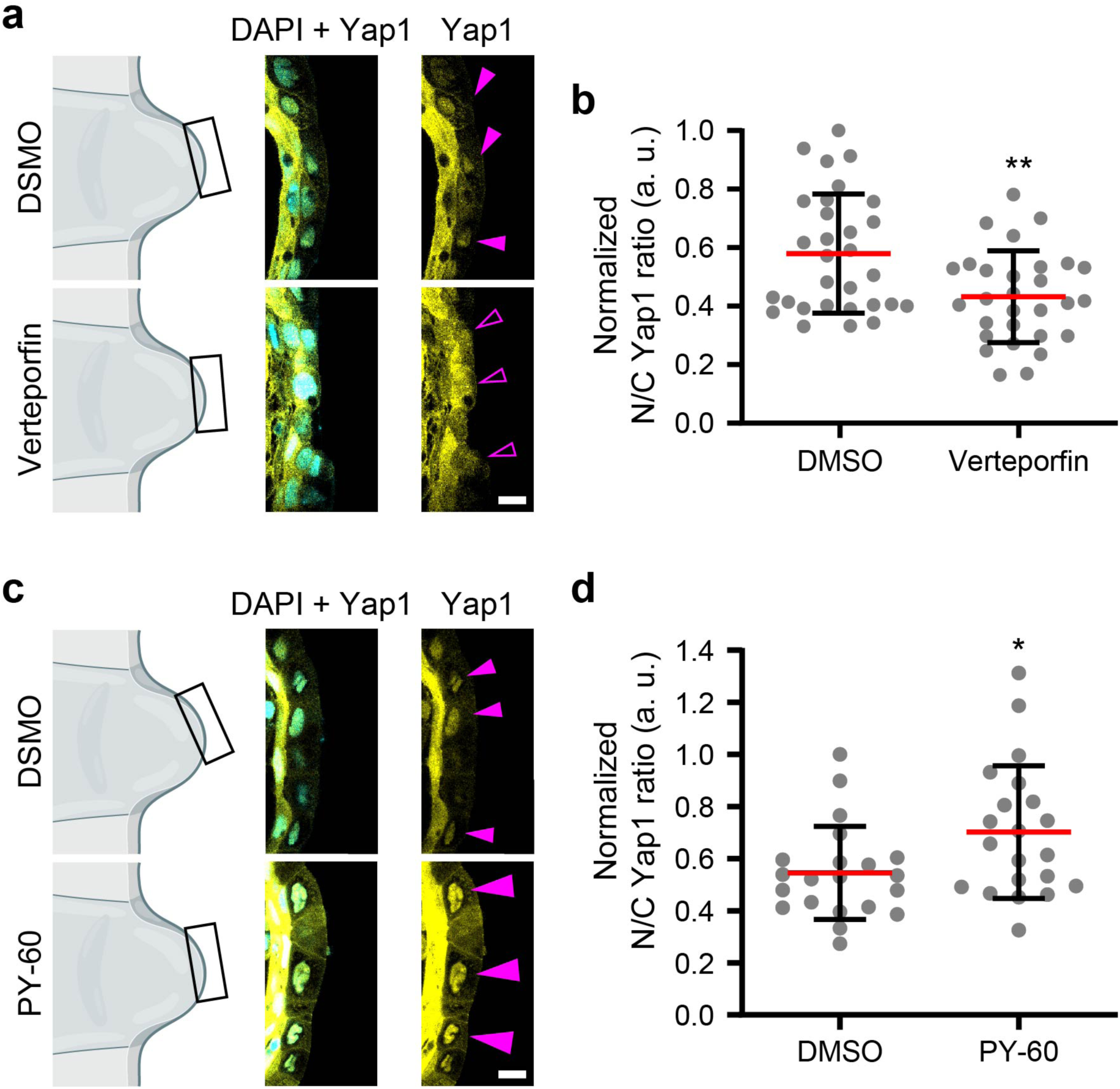
**Yap1 chemical inhibitor and activator functional controls. a,b**, Yap1 inhibitor Verteporfin reduces nuclear localization of Yap1 at the transition (stage NF41, 6 hpa). **a**, Left panels, schemes of the prospective regenerative bud with imaged area (black squares); Right panels, confocal slices of immunofluorescence against Yap1, with the nuclei counterstained with DAPI. Incubation with 0.01% DMSO or 2 µM Verteporfin. Filled magenta arrowheads, high nuclear localization of Yap1; outlined magenta arrowheads, low nuclear localization of Yap1. Scale bar, 30 µm. **b**, Quantification of Yap1 nucleo-cytoplasmic (N/C) ratio. Red lines represent mean and error bars standard deviation. Two-tailed Student’s *t-*test, \*\**p* = 0.0030, *n*_DMSO_ = 30, *n*_Verteporfin_ = 29 cells. **c,d**, Yap1 activator PY-60 augments nuclear localization of Yap1 at the transition (stage NF41, 6 hpa). **a**, Left panels, schemes of the prospective regenerative bud with imaged area (black squares); Right panels, confocal slices of immunofluorescence against Yap1, with the nuclei counterstained with DAPI. Incubation with 0.1% DMSO or 1 µM PY-60. Small magenta arrowheads, high nuclear localization of Yap1; Large magenta arrowheads, very high nuclear localization of Yap1. Scale bar, 30 µm. **d**, Quantification of Yap1 N/C ratio. Red lines represent mean and error bars standard deviation. Two-tailed Student’s *t*-test, \**p* = 0.0262, *n*_DMSO_ = 21, *n*_PY-60_ = 21 cells. **a,c**, Representative examples from at least three independent experiments; CI = 95%.

**Suppl Fig. 5.**
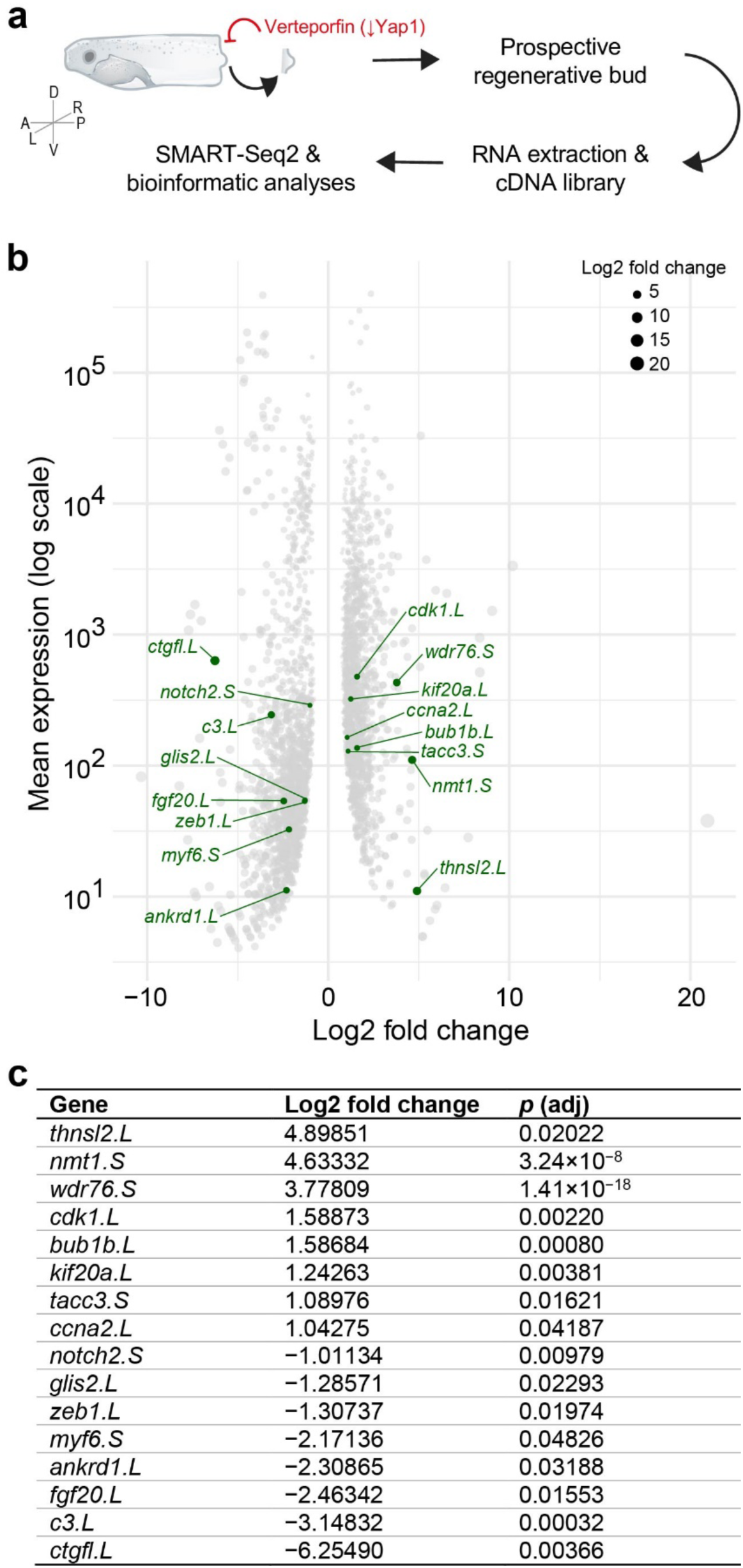
RNA-seq reveals a Yap1 transcriptional program at the transition from wound healing to regenerative bud formation. **a**, Scheme and flowchart for the collection and processing of isolated prospective regenerative buds at 6 hpa (stage NF41) for SMART-Seq2 RNA sequencing. A, anterior, P, posterior; D, dorsal; V, ventral; L, left; R, right. **b**, Volcano plot comparing Verteporfin (2 µM for 6 hpa) against regenerative wild-type with the expression and fold change of statistically differently expressing genes. Highlighted in green are genes known to be targets of Yap1. Positive values mean upregulated gene and negative values mean downregulated gene. **c**, Table with the numerical annotation of the highlighted genes in **b**.

**Suppl Fig. 6.**
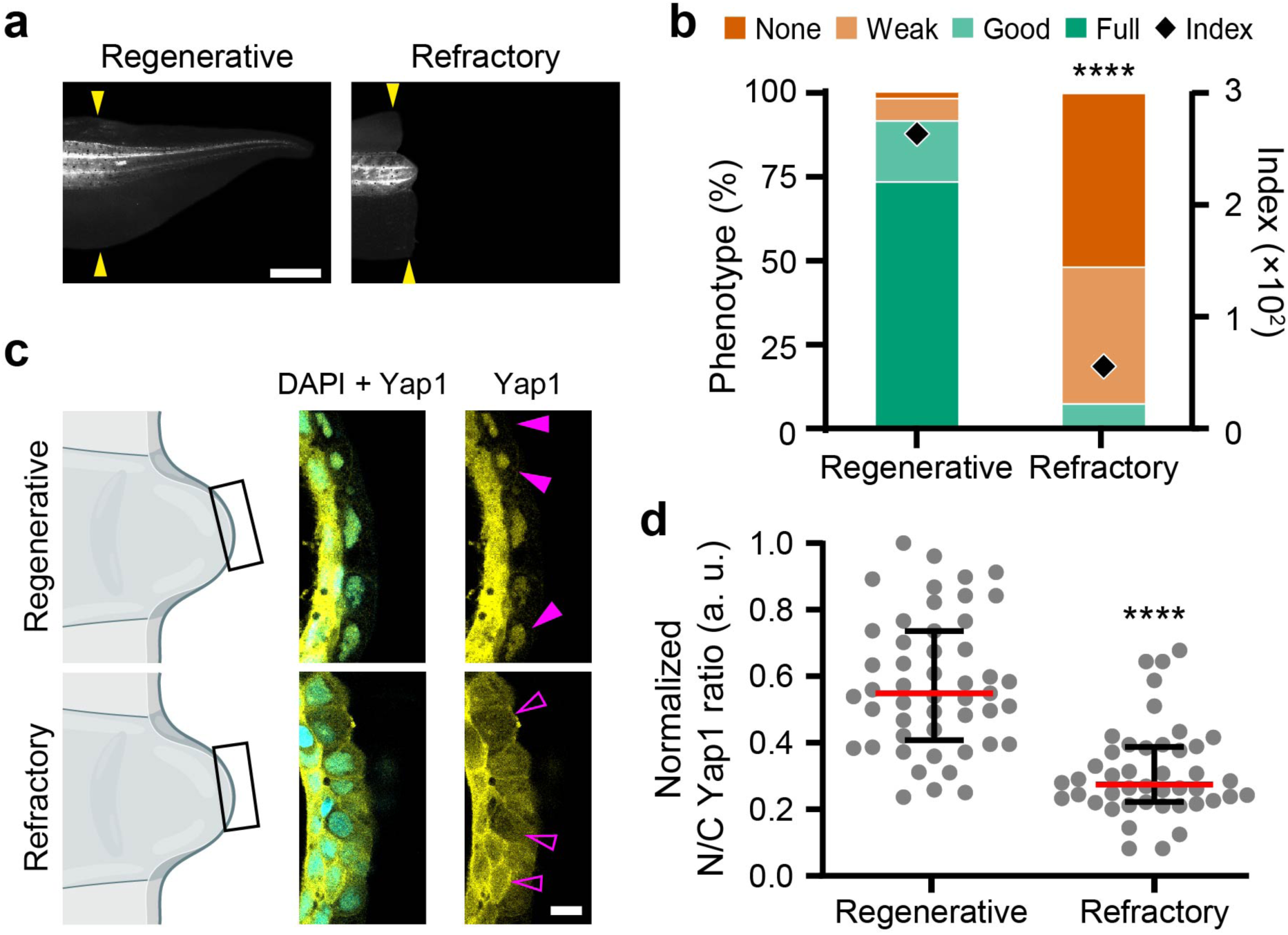
**Yap1 expression in nonregenerative refractory period (NF46) and regenerative stages (NF41). a,b**, Regeneration is abolished in the refractory period. **a**, Tails at 7 dpa from tadpoles amputated during the regenerative or refractory periods. Yellow arrowheads, amputation plane. Scale bar, 500 µm. **b**, Quantification of regeneration efficiency. Two-tailed Fisher’s exact test, *p* < 0.0001, *n*_Regenerative_ = 48, *n*_Refractory_ = 43 tails. **c,d**, Refractory period tails have reduced nuclear localization of Yap1 at the transition when compared to tails amputated in the regenerative period. **c**, Left panels, schemes of the prospective regenerative bud with imaged area (black squares); Right panels, confocal slices of immunofluorescence against Yap1, with the nuclei counterstained with DAPI. Incubation with 0.01% DMSO or 2 µM Verteporfin. Filled magenta arrowheads, high nuclear localization of Yap1; outlined magenta arrowheads, low nuclear localization of Yap1. Scale bar, 30 µm. **a,c**, Representative examples from at least three independent experiments; CI = 95%.

**Suppl Fig. 7.**
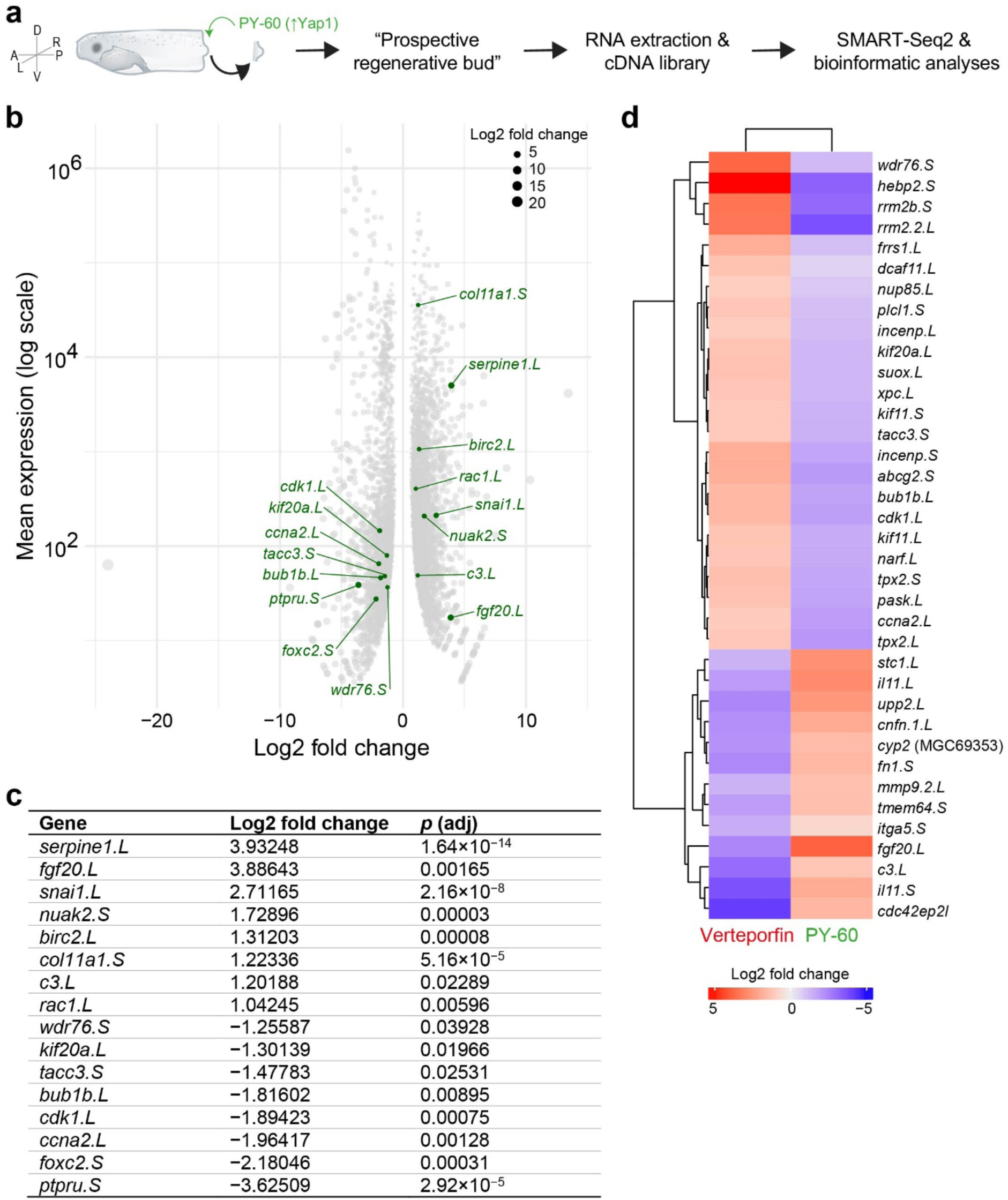
Pharmacological activation of Yap1 in the nonregenerative refractory period rescues the Yap1 transcriptional program as revealed by RNA-seq. **a**, Scheme and flowchart for the collection and processing of isolated tissues that topologically correspond to the “prospective regenerative bud” at 6 hpa in the refractory period (stage NF46) for SMART-Seq2 RNA sequencing. A, anterior, P, posterior; D, dorsal; V, ventral; L, left; R, right. **b**, Volcano plot comparing PY-60 (1 µM for 6 hpa) against refractory wild-type with the expression and fold change of statistically differently expressing genes. Highlighted in green are genes known to be targets of Yap1. Positive values mean upregulated gene and negative values mean downregulated gene. **c**, Table with the numerical annotation of the highlighted genes in **b**. **d**, Heatmap with fold change levels (extracted from **b** and **Suppl** Fig. 5b) of Yap1 target genes following pharmacological Yap1 inhibition or activation. Note that almost all these genes are known to be correlated with or required for vertebrate regeneration. Positive (red) values mean upregulated gene and negative (blue) values mean downregulated gene.

## References

1. Brockes, J. P. & Kumar, A. Comparative aspects of animal regeneration. Annu Rev Cell Dev Biol 24, 525–549 (2008).

2. Costa, E. C., Otsuki, L., Albors, A. R., Tanaka, E. M. & Chara, O. Spatiotemporal control of cell cycle acceleration during axolotl spinal cord regeneration. Elife 10, (2021).

3. Beck, C. W., Izpisúa Belmonte, J. C. & Christen, B. Beyond early development: Xenopus as an emerging model for the study of regenerative mechanisms. Dev Dyn 238, 1226–48 (2009).

4. Alvarado, A. S. & Tsonis, P. A. Bridging the regeneration gap: genetic insights from diverse animal models. Nat Rev Genet 7, 873–84 (2006).

5. Poss, K. D. & Tanaka, E. M. Hallmarks of regeneration. Cell Stem Cell (2024) doi:10.1016/J.STEM.2024.07.007.

6. Goldman, J. A. & Poss, K. D. Gene regulatory programmes of tissue regeneration. Nature Reviews Genetics vol. 21 511–525 Preprint at 10.1038/s41576-020-0239-7 (2020).

7. Tanaka, E. M. & Reddien, P. W. The Cellular Basis for Animal Regeneration. Dev Cell 21, 172– 185 (2011).

8. Chiou, K. & Collins, E. M. S. Why we need mechanics to understand animal regeneration. Dev Biol 433, 155–165 (2018).

9. Cochet-Escartin, O., Locke, T. T., Shi, W. H., Steele, R. E. & Collins, E.-M. S. Physical Mechanisms Driving Cell Sorting in Hydra. Biophys J 113, 2827–2841 (2017).

10. Huang, T.-Y. et al. Cooperative Regulation of Substrate Stiffness and Extracellular Matrix Proteins in Skin Wound Healing of Axolotls. Biomed Res Int 2015, 1–10 (2015).

11. Riquelme-Guzmán, C. et al. In vivo assessment of mechanical properties during axolotl development and regeneration using confocal Brillouin microscopy. Open Biol 12, (2022).

12. Edwards-Jorquera, S. et al. Mechanical control of tissue growth during limb regeneration. bioRxiv 2025.04.07.647008 (2025) doi:10.1101/2025.04.07.647008.

13. Mui, B. W. H. et al. Hyaluronic Acid and Emergent Tissue Mechanics Orchestrate Digit Tip Regeneration. bioRxiv 2024.12.04.626830 (2024) doi:10.1101/2024.12.04.626830.

14. Barriga, E. H., Franze, K., Charras, G. & Mayor, R. Tissue stiffening coordinates morphogenesis by triggering collective cell migration in vivo. Nature 554, 523–527 (2018).

15. Mao, Y. & Wickström, S. A. Mechanical state transitions in the regulation of tissue form and function. Nature Reviews Molecular Cell Biology 2024 1–17 (2024) doi:10.1038/s41580-024-00719-x.

16. Mongera, A. et al. A fluid-to-solid jamming transition underlies vertebrate body axis elongation. Nature 2018 561:7723 561, 401–405 (2018).

17. Shroff, N. P. et al. Proliferation-driven mechanical compression induces signalling centre formation during mammalian organ development. Nature Cell Biology 2024 26:4 26, 519–529 (2024).

18. Aztekin, C. Mechanisms of regeneration: to what extent do they recapitulate development? Development 151, (2024).

19. Prasad, K. & Palakodeti, D. Cellular and molecular mechanisms of development and regeneration. Development (Cambridge) 151, (2024).

20. Vining, K. H. & Mooney, D. J. Mechanical forces direct stem cell behaviour in development and regeneration. Nature Reviews Molecular Cell Biology 2017 18:12 18, 728–742 (2017).

21. Phipps, L. S., Marshall, L., Dorey, K. & Amaya, E. Model systems for regeneration: Xenopus. Development (Cambridge) 147, (2020).

22. Ferreira, F., Raghunathan, V., Luxardi, G., Zhu, K. & Zhao, M. Early redox activities modulate Xenopus tail regeneration. Nat Commun 9, 1–15 (2018).

23. Aztekin, C. et al. Identification of a regeneration-organizing cell in the Xenopus tail. Science (1979) 364, 653–658 (2019).

24. Sindelka, R. et al. Characterization of regeneration initiating cells during Xenopus laevis tail regeneration. Genome Biology 2024 25:1 25, 1–28 (2024).

25. Kim, H. Y., Jackson, T. R., Stuckenholz, C. & Davidson, L. A. Tissue mechanics drives regeneration of a mucociliated epidermis on the surface of Xenopus embryonic aggregates. Nat Commun 11, 665 (2020).

26. Koser, D. E. et al. Mechanosensing is critical for axon growth in the developing brain. Nat Neurosci 19, 1592–1598 (2016).

27. Rauzi, M., Verant, P., Lecuit, T. & Lenne, P. F. Nature and anisotropy of cortical forces orienting Drosophila tissue morphogenesis. Nat Cell Biol 10, 1401–1410 (2008).

28. Bjerke, M. A., Dzamba, B. J., Wang, C. & DeSimone, D. W. FAK is required for tension-dependent organization of collective cell movements in Xenopus mesendoderm. Dev Biol 394, 340–356 (2014).

29. Wang, N. et al. Mechanical behavior in living cells consistent with the tensegrity model. Proc Natl Acad Sci U S A 98, 7765–7770 (2001).

30. Pogoda, K. et al. Compression stiffening of brain and its effect on mechanosensing by glioma cells. New J Phys 16, 075002 (2014).

31. Storm, C., Pastore, J. J., MacKintosh, F. C., Lubensky, T. C. & Janmey, P. A. Nonlinear elasticity in biological gels. Nature 435, 191–194 (2005).

32. Chien, Y. H., Keller, R., Kintner, C. & Shook, D. R. Mechanical strain determines the axis of planar polarity in ciliated epithelia. Current Biology 25, 2774–2784 (2015).

33. Gudipaty, S. A. et al. Mechanical stretch triggers rapid epithelial cell division through Piezo1. Nature 543, 118–121 (2017).

34. Eisenhoffer, G. T. et al. Crowding induces live cell extrusion to maintain homeostatic cell numbers in epithelia. Nature 2012 484:7395 484, 546–549 (2012).

35. Tytti, K. et al. Mechanosensitive TRPV4 channel guides maturation and organization of the bilayered mammary epithelium. Scientific Reports 2024 14:1 14, 1–15 (2024).

36. Marchant, C. L., Malmi-Kakkada, A. N., Espina, J. A. & Barriga, E. H. Cell clusters softening triggers collective cell migration in vivo. Nature Materials 2022 1–10 (2022) doi:10.1038/s41563-022-01323-0.

37. Canales Coutiño, B. & Mayor, R. The mechanosensitive channel Piezo1 cooperates with semaphorins to control neural crest migration. Development 148, (2021).

38. Ferreira, F., Moreira, S., Zhao, M. & Barriga, E. H. Stretch-induced endogenous electric fields drive directed collective cell migration in vivo. Nature Materials 2025 1–9 (2025) doi:10.1038/s41563-024-02060-2.

39. Suchyna, T. M. et al. Identification of a peptide toxin from Grammostola spatulata spider venom that blocks cation-selective stretch-activated channels. Journal of General Physiology 115, 583–598 (2000).

40. Elosegui-Artola, A. et al. Force Triggers YAP Nuclear Entry by Regulating Transport across Nuclear Pores. Cell 171, 1397–1410.e14 (2017).

41. Dupont, S. et al. Role of YAP/TAZ in mechanotransduction. Nature 2011 474:7350 474, 179–183 (2011).

42. Zanconato, F. et al. Genome-wide association between YAP/TAZ/TEAD and AP-1 at enhancers drives oncogenic growth. Nature Cell Biology 2015 17:9 17, 1218–1227 (2015).

43. Pocaterra, A., Romani, P. & Dupont, S. YAP/TAZ functions and their regulation at a glance. J Cell Sci 133, (2020).

44. Hayashi, S. et al. Transcriptional regulators in the Hippo signaling pathway control organ growth in Xenopus tadpole tail regeneration. Dev Biol 396, 31–41 (2014).

45. Hayashi, S., Tamura, K. & Yokoyama, H. Yap1, transcription regulator in the Hippo signaling pathway, is required for Xenopus limb bud regeneration. Dev Biol 388, 57–67 (2014).

46. Li, J., Zhang, S. & Amaya, E. The cellular and molecular mechanisms of tissue repair and regeneration as revealed by studies in Xenopus. Regeneration (Oxf) 3, 198–208 (2016).

47. Wang, C. et al. Verteporfin inhibits YAP function through up-regulating 14-3-3σ sequestering YAP in the cytoplasm. Am J Cancer Res 6, 27–37 (2016).

48. Shalhout, S. Z. et al. YAP-dependent proliferation by a small molecule targeting annexin A2. Nat Chem Biol 17, 767–775 (2021).

49. Botello-Smith, W. M. et al. A mechanism for the activation of the mechanosensitive Piezo1 channel by the small molecule Yoda1. Nat Commun 10, 1–10 (2019).

50. Syeda, R. et al. Chemical activation of the mechanotransduction channel Piezo1. Elife 4, (2015).

51. Gregorieff, A., Liu, Y., Inanlou, M. R., Khomchuk, Y. & Wrana, J. L. Yap-dependent reprogramming of Lgr5+ stem cells drives intestinal regeneration and cancer. Nature 2015 526:7575 526, 715–718 (2015).

52. Kim, M., Kim, T., Johnson, R. L. & Lim, D. S. Transcriptional co-repressor function of the hippo pathway transducers YAP and TAZ. Cell Rep 11, 270–282 (2015).

53. Moya, I. M. & Halder, G. Hippo–YAP/TAZ signalling in organ regeneration and regenerative medicine. Nat Rev Mol Cell Biol 20, 211–226 (2019).

54. Zhong, Z., Jiao, Z. & Yu, F. X. The Hippo signaling pathway in development and regeneration. Cell Rep 43, (2024).

55. Basu, S., Totty, N. F., Irwin, M. S., Sudol, M. & Downward, J. Akt phosphorylates the Yes-associated protein, YAP, to induce interaction with 14-3-3 and attenuation of p73-mediated apoptosis. Mol Cell 11, 11–23 (2003).

56. Chow, C.-W. & Davis, R. J. Integration of Calcium and Cyclic AMP Signaling Pathways by 14-3-3. Mol Cell Biol 20, 702–712 (2000).

57. Patel, J. H., Schattinger, P. A., Takayoshi, E. E. & Wills, A. E. Hif1α and Wnt are required for posterior gene expression during Xenopus tropicalis tail regeneration. Dev Biol 483, 157–168 (2022).

58. Patel, J. H., Ong, D. J., Williams, C. R., Callies, L. L. K. & Wills, A. E. Elevated pentose phosphate pathway flux supports appendage regeneration. Cell Rep 41, 111552 (2022).

59. Lin, W. et al. Reactivation of mammalian regeneration by turning on an evolutionarily disabled genetic switch. Science (1979) 388, (2025).

60. Ransom, R. C. et al. Mechanoresponsive stem cells acquire neural crest fate in jaw regeneration. Nature 2018 563:7732 563, 514–521 (2018).

## References for Methods

61. Sive, H. L., Grainger, R. M. & Harland, R. M. Xenopus laevis In Vitro Fertilization and Natural Mating Methods. Cold Spring Harb Protoc (2007) doi:10.1101/pdb.prot4737.

62. Nieuwkoop, P. D. & Faber, J. Normal Table of Xenopus Laevis (Daudin). (Amsterdam: North-Holland, 1967).

63. Adams, D. S., Masi, A. & Levin, M. H+ pump-dependent changes in membrane voltage are an early mechanism necessary and sufficient to induce Xenopus tail regeneration. Development 134, 1323–35 (2007).

64. Love, N. R. et al. Amputation-induced reactive oxygen species are required for successful Xenopus tadpole tail regeneration. Nat Cell Biol 15, 222–8 (2013).

65. Bae, C., Sachs, F. & Gottlieb, P. A. The mechanosensitive ion channel Piezo1 is inhibited by the peptide GsMTx4. Biochemistry 50, 6295–6300 (2011).

66. Moreira, S., Espina, J. A., Saraiva, J. E. & Barriga, E. H. A Toolbox to Study Tissue Mechanics In Vivo and Ex Vivo. Methods in Molecular Biology 2438, 495–515 (2022).

67. Hutter, J. L. & Bechhoefer, J. Calibration of atomic-force microscope tips. Review of Scientific Instruments 64, 1868–1873 (1993).

68. Hertz, H. Über die Berührung fester elastischer Körper. J reine und angewandte Mathematik 92, 156–171 (1881).

69. Das, A., Fischer, R. S., Pan, D. & Waterman, C. M. YAP Nuclear Localization in the Absence of Cell-Cell Contact Is Mediated by a Filamentous Actin-dependent, Myosin II-and Phospho-YAP-independent Pathway during Extracellular Matrix Mechanosensing. Journal of Biological Chemistry 291, 6096–6110 (2016).

70. Macaulay, I. C. et al. Separation and parallel sequencing of the genomes and transcriptomes of single cells using G&T-seq. Nat Protoc 11, 2081–2103 (2016).

71. Baym, M. et al. Inexpensive Multiplexed Library Preparation for Megabase-Sized Genomes. PLoS One 10, e0128036 (2015).

72. Chen, S., Zhou, Y., Chen, Y. & Gu, J. fastp: an ultra-fast all-in-one FASTQ preprocessor. Bioinformatics 34, i884–i890 (2018).

73. Kim, D., Langmead, B. & Salzberg, S. L. HISAT: a fast spliced aligner with low memory requirements. Nature Methods 2015 12:4 12, 357–360 (2015).

74. Li, H. et al. The Sequence Alignment/Map format and SAMtools. Bioinformatics 25, 2078–2079 (2009).

75. Liao, Y., Smyth, G. K. & Shi, W. featureCounts: an efficient general purpose program for assigning sequence reads to genomic features. Bioinformatics 30, 923–930 (2014).

76. Love, M. I., Huber, W. & Anders, S. Moderated estimation of fold change and dispersion for RNA-seq data with DESeq2. Genome Biol 15, 1–21 (2014).

77. Gu, Z., Eils, R. & Schlesner, M. Complex heatmaps reveal patterns and correlations in multidimensional genomic data. Bioinformatics 32, 2847–2849 (2016).

